# A minimum module for positioning the Chromosomal Passenger Complex at the cell center for cytokinesis

**DOI:** 10.1101/2025.09.02.673761

**Authors:** Konstantinos Nakos, Sithara Wijeratne, Michael Blower, Radhika Subramanian

## Abstract

The Aurora B kinase-containing Chromosomal Passenger Complex (CPC) is an essential regulator of cytokinesis, the final and irreversible step of cell division. During anaphase, the CPC concentrates at the spindle midzone - a bundle of overlapping antiparallel microtubules organized at the cell center. How CPC is selectively enriched at the midzone within the dense, heterogeneous, and dynamic spindle microtubule network is unknown. Here, we define a minimal CPC midzone enrichment module. We show that the maximal enrichment of CPC at antiparallel microtubule overlaps requires PRC1-crosslinked microtubules and the interaction of CPC with two mitotic kinesins, KIF4A and KIF20A. We find that the two motors exhibit a division of labor: KIF20A delivers CPC from non-overlapping microtubules, and KIF4A retains CPC at PRC1-crosslinked overlaps. Conditional depletion of KIF4A in mitotic cells reveals that it is required for CPC localization at the spindle midzone in anaphase. Taken together, our findings reveal how the collective activity of two kinesins enables navigation through the complex microtubule network of the spindle to organize kinase signaling at the cell center for cytokinesis.

## INTRODUCTION

Genome integrity relies on the precisely coordinated spatial and temporal regulation of mitotic kinases during cell division. Mitotic progression is often accompanied by the dynamic repositioning of kinases to specific sites within the spindle apparatus. A striking example is the essential Aurora B kinase, a component of the evolutionarily conserved Chromosomal Passenger Complex (CPC), named for its dynamic localization and activity at distinct subcellular sites throughout mitosis^1^. At the metaphase-to-anaphase transition, the CPC relocates from centromeres to the spindle midzone^2,3^, where it phosphorylates effectors such as MKLP1, KIF4A, and MgcRacGAP1 to ensure error-free cytokinesis -the final and irreversible step of mitosis^4–8^.

The spindle midzone is a specialized array of overlapping antiparallel microtubules that assembles at the cell center during anaphase. It functions as a signaling platform for the recruitment of key cytokinesis factors, including the CPC^9,10^. In metazoan cells, the midzone forms through crosslinking of microtubules from opposite spindle halves at anaphase onset. As anaphase progresses, sliding of microtubules by motor proteins causes the overlap region to narrow until it reaches a stable length of ∼2–3 µm, further concentrating the midzone-associated proteins at the cell center^10–13^. *In vitro* studies have shown that coordinated “bundling, sliding, and compaction” of microtubules by the PRC1-KIF4A module can recapitulate midzone organization^14–16^. PRC1 (Protein Regulator of Cytokinesis-1) is a non-motor protein that preferentially crosslinks and accumulates at the overlap region of antiparallel microtubules^17–19^. KIF4A is a plus-end-directed kinesin that is recruited to these overlaps via a high-affinity interaction with PRC1, where it controls overlap length^17,18^. KIF4A suppresses tubulin polymerization at microtubule plus-ends, and crosslinks and slides microtubules apart^14–16,20,21^. This narrows the antiparallel overlap and increases local protein density, eventually stalling further microtubule sliding and establishing a stable overlap region^14,16^. A central question is how the CPC is specifically delivered to and retained at the narrow and dynamically organized subset of midzone microtubules, within the larger and heterogeneous spindle network^17^.

CPC is a microtubule-binding protein complex, and this activity is required for its midzone localization^22–24^. The complex is organized around the scaffold protein INCENP, with its N-terminus binding Borealin and Survivin to form the centromere-targeting module, and its C-terminal IN-box associating with Aurora B to form the active kinase (Figure 1A)^2,3^. These domains are connected by a poorly characterized >600-amino acid segment referred to as the INCENP tether, which contains an Intrinsically Disordered Region (IDR: aa101–527) and a Single Alpha Helix (SAH: aa539–747)^23,25,26^. The SAH domain is proposed to serve as a flexible spacer between the centromere-targeting and kinase modules^1,23,26^. This domain also contains the microtubule-binding site (hereafter “SAH-MT”) required for midzone localization (Figure 1A)^23,24,27^. Borealin harbors a second microtubule-binding domain in CPC^28^. *In vitro*, *Xenopus laevis* CPC and soluble tubulin heterodimers form condensates that result in the assembly of parallel microtubule bundles^29^. Whether CPC’s intrinsic microtubule-binding and bundling activities contribute to its enrichment at antiparallel microtubule overlaps is not known.

**Figure 1:**
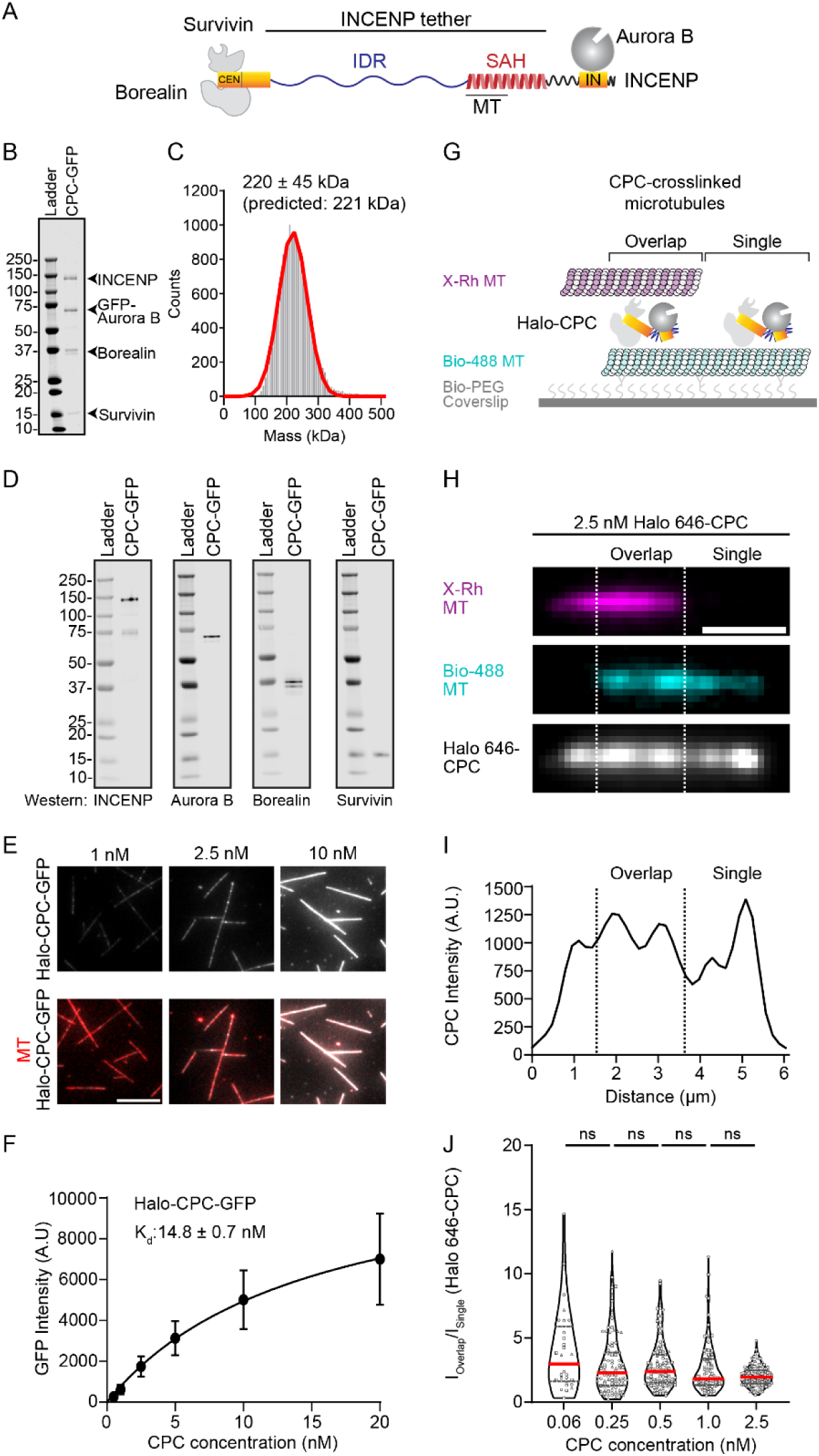
CPC microtubule-binding and crosslinking activity are insufficient for enrichment at microtubule overlaps. (A) The cartoon shows human CPC, which consists of INCENP, Aurora B, Borealin, and Survivin. The INCENP tether consists of IDR (Intrinsically Disordered Region) and SAH (Single alpha helix). IN; IN-box, CEN; Centromeric binding region, SAH-MT; Microtubule-binding domain. (B) SDS-PAGE of purified human CPC-GFP. Arrowheads show INCENP, GFP-Aurora B, Borealin, and Survivin. (C) Mass photometry of CPC-GFP. (D) Western blots of individual subunits of CPC-GFP using antibodies specific for INCENP, Aurora B, Borealin, and Survivin. (E) Representative fluorescent images of Halo-CPC-GFP bound to single immobilized microtubules at the indicated concentrations. Top row panels show CPC (grey), and bottom row panels are merged images of CPC (grey) and microtubules (red). The scale bar is 10 μm. (F) Binding curve of the interaction between Halo-CPC-GFP and microtubules. Data is represented as mean ± SD. Data were fit to a Hill equation with K_d_: 14.8 ± 0.7 nM. 0.5 nM N=329 (mean ± SD: 262.1 ± 100.3), 1 nM N=383 (mean ± SD: 618.9 ± 230.3), 2.5 nM N=371 (mean ± SD: 1748 ± 503.4), 5 nM N=412 (mean ± SD: 3135 ± 839.6), 10 nM N=397 (mean ± SD: 5018 ±1441), 20 nM N=336 (mean ± SD: 7000 ± 2243). (G) Schematic of the *in vitro* TIRFM assay to examine the enrichment of Halo 646-CPC at CPC-crosslinked microtubule overlaps. (H) The representative fluorescent images show the localization of Halo 646-CPC (2.5 nM, grey) at a microtubule overlap (magenta and cyan; region between the dotted lines). The scale bar is 2 μm. (I) The graph shows the intensity profile of Halo 646-CPC in H. (J) Violin plot of Halo 646-CPC I_Overlap_/I_Single_ ratio at different CPC concentrations. Horizontal lines indicate the median (red line) and 25^th^ and 75^th^ percentiles (grey dotted lines). 0.06 nM: N=29 (median [IQR]: 2.97 [1.63,5.89]), 0.25 nM: N=95 (median [IQR]: 2.28 [1.29,3.87]), 0.5 nM: N=103 (median [IQR]: 2.39 [1.51, 3.7]), 1 nM: N=96 (median [IQR]: 1.82 [1.34,3.36]) and 2.5 nM: N=137 (median [IQR]: 1.96 [1.46, 2.49]). ns, not significant (p>0.05).

In addition to its intrinsic microtubule-bundling activity, current models of CPC midzone localization are also based on its interaction with the kinesin-6 motor KIF20A (“CPC-KIF20A” module; human KIF20A is also referred to as MKLP2). In the absence of KIF20A, CPC fails to dissociate from chromosomes^30–34^. *In vitro*, truncated CPC constructs lacking the SAH-MT domain can be transported by KIF20A to the plus-ends of individual microtubules^34^. KIF20A also exhibits microtubule-crosslinking activity^32,35^. Thus, KIF20A-mediated CPC transport and microtubule crosslinking are thought to be the mechanism by which the KIF20A-CPC module enriches CPC at antiparallel microtubule overlaps. However, this protein module has not been reconstituted in the context of crosslinked microtubules, and whether it is sufficient to enrich CPC at overlaps is unknown.

Our current understanding of midzone organization in human cells involves distinct functional protein modules with an architectural role for the PRC1-KIF4A module and a CPC-enrichment role for the CPC-KIF20A module. How the activities of the two modules are coordinated for CPC midzone enrichment remains unclear. Depletion of PRC1 and KIF4A in human cells leads to changes in the localization of most midzone proteins, including the CPC^36–38^. However, CPC mislocalization under these conditions is largely attributed to indirect effects of perturbing the midzone microtubule organization and not due to direct binding to CPC. Many midzone proteins, including PRC1 and KIF4A, are reported in a proteomics study of Aurora B kinase interactors in HeLa cells. However, whether these represent direct binding or indirect associations through other midzone proteins is not known^39^. KIF4A is an Aurora B substrate, and its phosphorylation enhances its ability to regulate microtubule length, but whether there are additional interactions between CPC and KIF4A is not known^5,39^. In *Xenopus* egg extracts, the KIF4A homolog (XKLP1) contributes to CPC transport along astral microtubules to the interaction zone of large (∼100 mm) asters^40^. However, in human somatic cells, KIF4A is mainly confined to the PRC1-crosslinked midzone. In general, dissecting the contributions of each component of the two modules to CPC enrichment at the midzone has been challenging in cell biological studies, as each protein plays essential and sometimes overlapping roles in metaphase function or midzone organization, or both.

*In vitro* reconstitution approaches are powerful in elucidating the minimal set of activities required to recapitulate a cellular phenomenon in a cell-free system. However, reconstitutions of CPC midzone targeting have been limited by technical challenges in purifying human CPC with an intact INCENP tether that retains complete microtubule-binding activity. Here, we overcome this barrier by generating a soluble, full-length recombinant human CPC (“holo-CPC”). Using a systematic bottom-up reconstitution approach, we dissected the individual and combined contributions of three midzone microtubule crosslinking proteins (PRC1, KIF4A, KIF20A) and CPC itself to the selective enrichment of CPC at overlap regions of PRC1-crosslinked microtubules. To our surprise, the CPC-KIF20A module was not sufficient to concentrate CPC at these overlaps. Instead, we found that CPC is directly transported by KIF4A, which is important for CPC localization at microtubule overlaps *in vitro* and at the midzone in human cells. Addition of KIF20A to the reconstituted PRC1-KIF4A-CPC network further enhanced CPC enrichment at antiparallel overlaps. Together, our results define a minimal “CPC midzone enrichment module” and uncover the mechanism by which the collective activity of two motors specifically concentrates the CPC at the overlap region of PRC1-crosslinked antiparallel microtubules.

## RESULTS

### *In vitro* reconstitution of human CPC reveals that intrinsic microtubule binding and bundling activities do not enrich CPC at microtubule overlaps

We co-expressed full-length human Borealin, Survivin, INCENP, and GFP-tagged Aurora B in insect cells and established a protocol to purify recombinant GFP-tagged holo-CPC (hereafter CPC-GFP) (Methods). SDS-PAGE and Western blot analysis from the peak fraction from size exclusion chromatography confirmed the presence of all four protein components in the purified complex (Figures 1B-D, S1A). Mass photometry revealed a single peak with a molecular weight of 220 ± 45 kDa, in agreement with the predicted molecular weight of 221 kDa (Figure 1C). These data confirm that the purified human CPC is a heterotetramer with one copy of each subunit.

Prior studies have shown that the microtubule interaction of CPC is required for its midzone localization^22–24^. We characterized the microtubule binding activity of the recombinant human CPC using a multi-wavelength Total Internal Reflection Fluorescence Microscopy (TIRFM) assay. Taxol-stabilized GMPCPP microtubules, labeled with X-Rhodamine and biotin, were immobilized on a neutravidin-functionalized glass coverslip and incubated with increasing concentrations of purified Halo-CPC-GFP. CPC molecules bound uniformly along microtubules with an apparent binding affinity of 14.8 ± 0.7 nM (Figures 1E-F, S1B-D).

We next examined if CPC could crosslink microtubules and whether it was preferentially enriched at regions of microtubule overlaps using a TIRFM-based microtubule bundling assay^14,20,41^. Biotinylated taxol-stabilized GMPCPP microtubules labeled with HiLyte 488 were surface immobilized and incubated with purified JFX646-conjugated Halo-CPC (Halo 646-CPC) (Figure S1E). Subsequently, X-Rhodamine-labeled, non-biotinylated microtubules were added to form CPC-crosslinked microtubule bundles. After washing away unbound microtubules and proteins, additional Halo 646-CPC, at the indicated concentrations, was flowed into the chamber. The binding reaction was given time to reach steady-state, and imaged. Microtubule overlap regions were identified by merging the 488 and X-Rhodamine-labeled microtubule images (Figure 1G). To quantify enrichment, we calculated the ratio of fluorescence intensity at overlaps relative to single microtubules (I_Overlap_/I_Single_). Since the maximum expected occupancy at an overlap formed by two microtubules is twice the occupancy at single microtubules, values of I_Overlap_/I_Single_ greater than 2 indicate specific enrichment at microtubule overlaps. For PRC1 homologs, which are the best studied midzone-specific MAPs, the reported I_Overlap_/I_Single_ ratio is ∼10 in similar *in vitro* assays^15,41^.

Our observations confirmed that CPC crosslinks microtubules *in vitro* (Figure 1H)^28,29^. However, as shown in the representative images and corresponding fluorescence line scan, CPC did not exhibit substantial enrichment at microtubule overlaps compared to single microtubules (Figures 1H-I). Ratiometric intensity analysis (I_Overlap_/I_Single_) showed approximately 1.8-3-fold enrichment of CPC at microtubule overlaps relative to single microtubules across all tested concentrations (Figure 1J). At the lowest CPC concentrations (0.06 nM and 0.25 nM), the intensity ratio had a broad distribution, due to sparse CPC binding, and dominant contribution from occasional CPC clusters bound to microtubules (Figure S1F-G).

We report the successful purification of soluble recombinant human holo-CPC. Characterization of the CPC-microtubule interactions reveals that while CPC has intrinsic microtubule-binding and crosslinking activity, this property alone is insufficient for selective enrichment of CPC at microtubule overlaps.

### The midzone proteins KIF20A and PRC1 do not enrich CPC at microtubule overlaps

We next examined the kinesin KIF20A, a known CPC-interacting motor protein, as a candidate factor for concentrating CPC at microtubule overlap regions^30–34^. Given that KIF20A possesses intrinsic microtubule crosslinking activity^32,35^, we first examined whether KIF20A intrinsically prefers to localize at microtubule overlaps. Using the protocol described in Figure 1G, we generated microtubule bundles with recombinant KIF20A-GFP (Figures 2A, S2A-B). To minimize microtubule sliding, which resulted in the separation of bundles at higher ATP concentrations, the assay was performed under lower ATP conditions (100 μM). Under these conditions, stable microtubule bundles were readily assembled across a broad range of KIF20A-GFP concentrations (Figure 2B). KIF20A-GFP was detected on both single microtubules and overlap regions (Figure 2B-C), exhibiting a modest enrichment of ∼2.2-3-fold at overlaps, with a slight dependence on protein concentration (Figure 2D).

**Figure 2:**
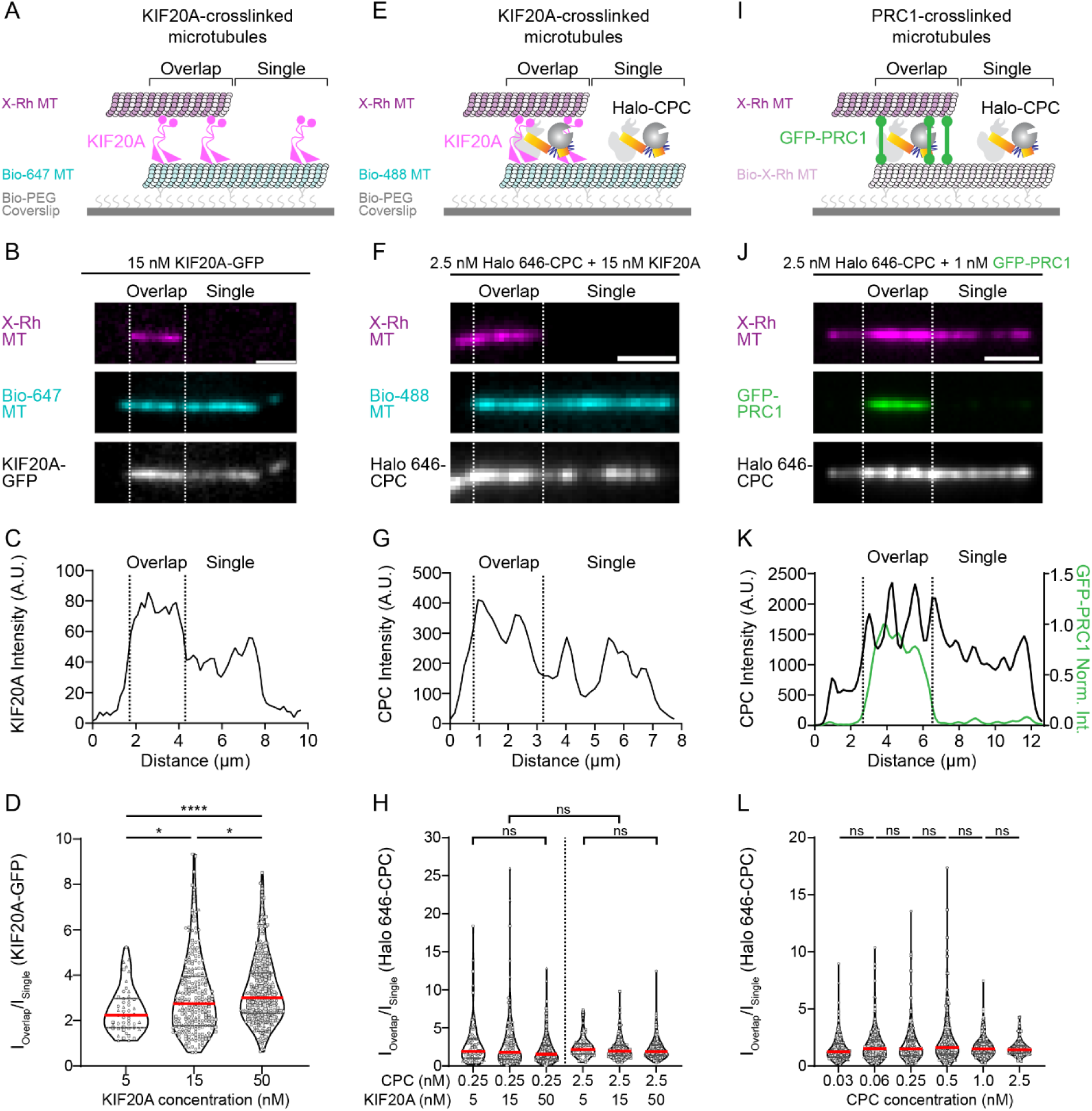
CPC does not enrich at microtubule overlaps formed by KIF20A or PRC1. (A) Schematic of the *in vitro* TIRFM assay to examine the enrichment of KIF20A-GFP at KIF20A-crosslinked microtubule overlaps. (B) The representative fluorescent images show the localization of KIF20A-GFP (15 nM, grey) on KIF20A-crosslinked microtubules (magenta and cyan; region between the dotted lines). The scale bar is 2 μm. (C) The graph shows the intensity profile of KIF20A-GFP in B. (D) Violin plot of KIF20A-GFP I_Overlap_/I_Single_ ratio at different KIF20A concentrations. Horizontal lines indicate the median (red line) and 25^th^ and 75^th^ percentiles (grey dotted lines). 5 nM KIF20A: N=51 (median [IQR]: 2.24 [1.68,2.96]), 15 nM KIF20A: N=239 (median [IQR]: 2.74 [1.78,3.93]), and 50 nM KIF20A: N=319 (median [IQR]: 3.00 [2.31,4.10]). ns, not significant (p>0.05). *p < 0.05, ****p< 0.0001. (E) Schematic of the *in vitro* TIRFM assay to examine the enrichment of Halo 646-CPC at KIF20A-crosslinked microtubule overlaps. (F) The representative fluorescent images show the localization of Halo 646-CPC (2.5 nM, grey) at a microtubule overlap (magenta and cyan; region between the dotted lines). The scale bar is 2 μm. (G) The graph shows the intensity profile of Halo 646-CPC in F. (H) Violin plot of Halo 646-CPC I_Overlap_/I_Single_ ratio at different CPC and KIF20A concentrations. Horizontal lines indicate the median (red line) and 25^th^ and 75^th^ percentiles (grey dotted lines). 0.25 nM Halo 646-CPC/5 nM KIF20A: N=67 (median [IQR]: 1.9 [1.02,3.53]), 0.25 nM Halo 646-CPC/15 nM KIF20A: N=118 (median [IQR]: 1.79 [0.86,3.74]), 0.25 nM Halo 646-CPC/50 nM KIF20A: N=206 (median [IQR]: 1.55 [0.88,2.69]), 2.5 nM Halo 646-CPC/5 nM KIF20A: N=70 (median [IQR]: 2.17 [1.56,2.94]), 2.5 nM Halo 646-CPC/15 nM KIF20A: N=165 (median [IQR]: 1.93 [1.34,2.69]) and 2.5 nM Halo 646-CPC/50 nM KIF20A: N=225 (median [IQR]: 1.88 [1.35,2.79]). ns, not significant (p>0.05). (I) Schematic of the *in vitro* TIRFM assay used to examine the enrichment of Halo 646-CPC at PRC1-crosslinked microtubule overlaps. (J) The representative fluorescent images show the localization of GFP-PRC1 (1 nM, green) and Halo 646-CPC (2.5 nM, grey) at a PRC1-crosslinked microtubule overlap (magenta; region between the dotted lines). The scale bar is 3 μm. (K) The graph shows the intensity profile of Halo 646-CPC and GFP-PRC1 in J. (L) Violin plot of Halo 646-CPC I_Overlap_/I_Single_ ratio at different CPC concentrations. Horizontal lines indicate the median (red line) and 25^th^ and 75^th^ percentiles (grey dotted lines). 0.03 nM Halo 646-CPC: N=179 (median [IQR]: 1.24 [0.78,1.95]), 0.06 nM Halo 646-CPC: N=206 (median [IQR]: 1.5 [0.81,2.3]), 0.25 nM Halo 646-CPC: N=188 (median [IQR]: 1.49 [0.9,2.27]), 0.5 nM Halo 646-CPC: N=208 (median [IQR]: 1.6 [1.05,2.65]), 1 nM Halo 646-CPC: N=189 (median [IQR]: 1.49 [1.08,2.03]) and 2.5 nM Halo 646-CPC: N=147 (median [IQR]: 1.43 [1.13,1.77]). ns, not significant (p>0.05).

We next tested whether KIF20A and CPC collectively promote CPC enrichment at microtubule overlaps. First, we verified that recombinant CPC binds KIF20A under TIRFM buffer conditions in the absence of microtubules. Pull-down assays confirmed the interaction between CPC and KIF20A (Figure S2C)^30–34^. We then examined Halo 646-CPC (0.25 nM and 2.5 nM) localization on KIF20A-crosslinked microtubule bundles assembled at varying KIF20A concentrations (Figure 2E). The assay was performed in the presence of 1 mM ATP to permit KIF20A-driven transport and microtubule sliding. Under conditions tested, CPC did not exhibit significant enrichment at overlap regions (Figure 2F-G). Consistent with visual observations, ratiometric intensity analysis (I_Overlap_/I_Single_) from steady-state images revealed 1.8-2.2-fold CPC enrichment at microtubule overlaps relative to single microtubules (Figure 2H). Notably, the enrichment of CPC at KIF20A-crosslinked microtubule overlaps was comparable to the enrichment seen in the absence of KIF20A (Figure 1H-J) and did not increase with higher concentrations of KIF20A (Figure 2H). These data indicate that CPC-KIF20A interactions alone are insufficient to drive CPC accumulation at overlap regions.

Finally, we examined PRC1, which selectively crosslinks and enriches at antiparallel overlaps, as a potential candidate for promoting CPC localization at microtubule overlaps^14,15,20,41^. PRC1-crosslinked overlaps were assembled with recombinant GFP-PRC1 (1 nM) and analyzed for Halo 646-CPC binding (Figures 2I, S2A-B). As expected, GFP-PRC1 displayed significant enrichment (∼10-fold) at microtubule overlaps (Figure 2J-K). In contrast, Halo 646-CPC exhibited only modest enrichment (∼1.2–1.6-fold) within the same bundles, across a range of CPC concentrations (Figure 2L). Thus, neither PRC1 nor antiparallel overlap geometry is sufficient to drive CPC accumulation at overlap regions.

Thus, none of the microtubule crosslinking midzone proteins tested (CPC, PRC1, and KIF20A) were sufficient to recapitulate the selective enrichment of CPC at microtubule overlaps, as observed in the spindle midzone.

### The PRC1-KIF4A-CPC-KIF20A network is the minimal module for CPC recruitment to antiparallel microtubule overlaps

Our experiments thus far did not recapitulate robust CPC enrichment at microtubule overlaps. We therefore turned to the midzone kinesin KIF4A, a known PRC1-binding protein^15,37,42,43^. Preassembled GFP-PRC1-crosslinked X-Rhodamine microtubule overlaps were incubated with 30 pM Halo 646-CPC in the presence of 15 nM KIF4A (Figures 3A-B, S3A-B). Strikingly, the presence of KIF4A led to a pronounced increase in Halo 646-CPC localization at PRC1-crosslinked overlaps (Figure 3B). Quantitative ratiometric analysis (I_Ioverlap_/I_Single_) revealed∼5-fold enrichment of CPC at overlaps relative to single microtubules, which is greater than any of the other conditions tested so far (Figure 3B, 3F). These results identify KIF4A as a molecular link between CPC and PRC1-crosslinked microtubule overlaps.

**Figure 3:**
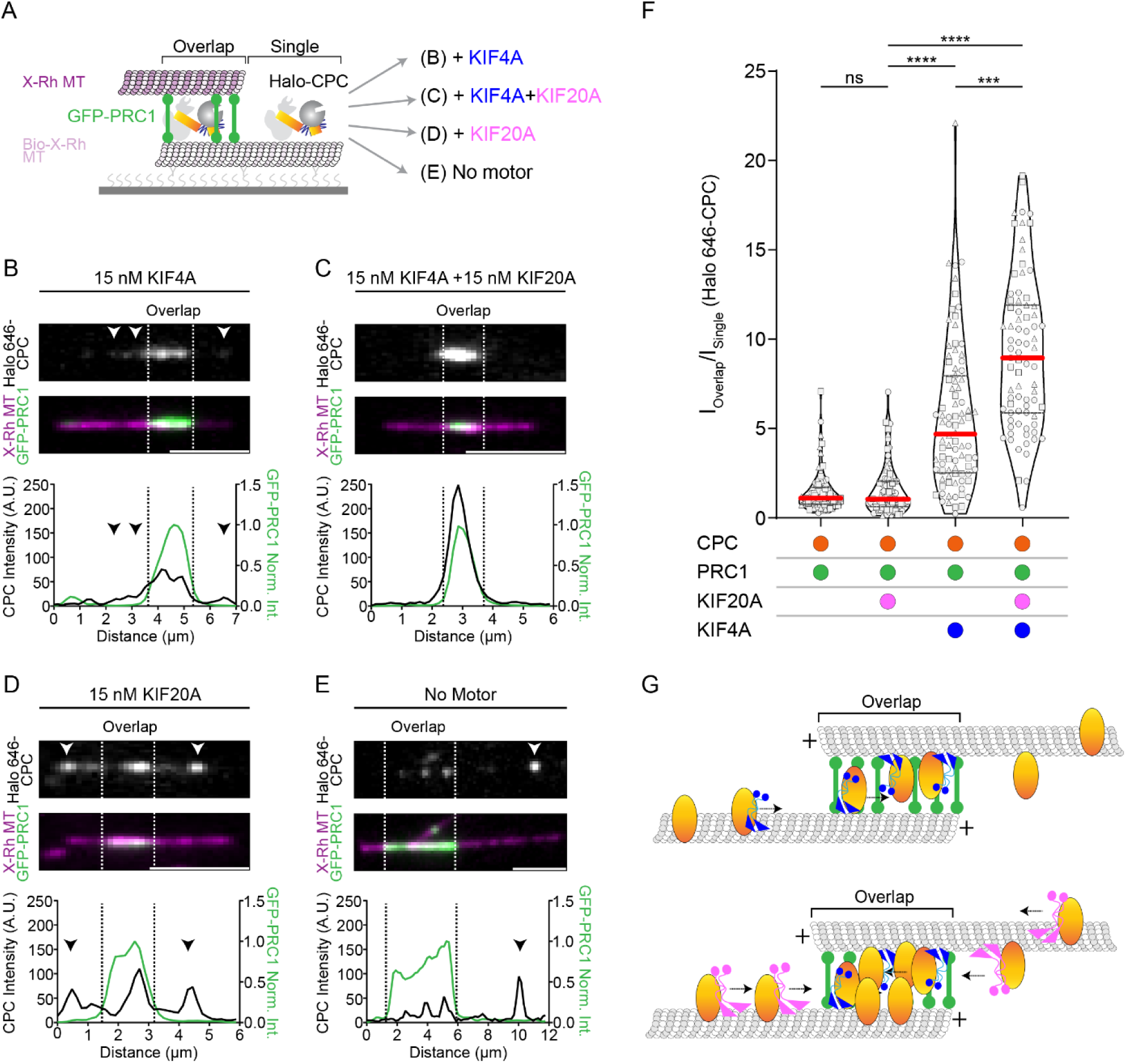
Reconstitution of a minimal module for CPC enrichment at PRC1-crosslinked microtubule overlaps. (A) Schematic of the *in vitro* TIRFM assay. (B-E) The fluorescent images show the localization of Halo 646-CPC (30 pM, grey) at crosslinked microtubule overlaps (magenta, regions between the dotted lines) formed by GFP-PRC1 (1 nM, green), in the presence of (B) KIF4A (15 nM), (C) KIF4A and KIF20A (15 nM each motor), (D) KIF20A (15 nM) or (E) in the absence of motor proteins. Arrowheads show Halo 646-CPC on single microtubules flanking the overlap. The graphs below show the fluorescence intensity profiles of Halo 646-CPC and GFP-PRC1 along the microtubule bundles. The scale bars are 3 μm. (F) Violin plots of Halo 646-CPC I_Overlap_/I_Single_ ratio from experiments in (B-E). Horizontal lines indicate the median (red line) and 25^th^ and 75^th^ percentiles (grey dotted lines). All experiments were at 30 pM Halo 646-CPC, 1nM GFP-PRC1, and 15 nM motors. CPC+PRC1: N=77 (median [IQR]: 1.11 [0.76,1.69]), CPC+PRC1+KIF20A: N=87 (median [IQR]: 1.04 [0.63,2.04]), CPC+PRC1+KIF4A: N=88 (median [IQR]: 4.68 [2.52,7.95]), and CPC+PRC1+KIF4A+KIF20A: N=79 (median [IQR]: 8.94 [5.88,11.9]). ns, nonsignificant (p > 0.05), ***p < 0.001, ****p < 0.0001. (G) The schematics illustrate the role of KIF4A (blue) as the CPC anchor at PRC1 (green)-crosslinked antiparallel microtubule overlap (top) and the synergistic activity of KIF20A (magenta) and KIF4A (blue) in maximizing the enrichment of CPC at overlaps relative to single microtubules.

Since CPC interacts with KIF20A, we next tested whether inclusion of KIF20A further impacts CPC recruitment to PRC1-crosslinked overlaps by KIF4A. Therefore, we repeated the reconstitution assay in the presence of 15 nM KIF4A and 15 nM KIF20A. Strikingly, the combined presence of both motor proteins resulted in ∼9-fold enrichment of CPC at PRC1-crosslinked microtubule overlaps (Figure 3C, 3F). For comparison, parallel reactions were performed either with KIF20A alone or in the absence of motor proteins, while maintaining constant concentrations of PRC1 and CPC (Figures 3D-E). In both control conditions, Halo 646-CPC localized to both overlaps and single microtubules, yielding I_Overlap_/I_Single_ ratios of ∼1 and ∼1.1, respectively (Figure 3D-F). Thus, KIF4A results in a ∼10-fold increase in CPC at PRC1-crosslinked overlaps compared to KIF20A in experiments with just one of the two motors (Figure 3B, 3D, 3F). In addition, these data demonstrate that KIF4A and KIF20A act synergistically to promote CPC accumulation at PRC1-crosslinked microtubule overlaps.

To gain further insight into the mechanism driving the enrichment of CPC at microtubule overlaps, we performed time-lapse imaging (Figure S3C, Movie 1). For these assays, Halo 646-CPC was used at 2.5 nM to enhance visualization of dynamic localization and accumulation. We first performed reactions with KIF4A (15 nM) as the sole motor protein. Consistent with previous reports, KIF4A recruitment by PRC1 induced relative sliding of the PRC1-crosslinked microtubule (Figure S3C, Movie 1A)^14,16^. Initially, Halo 646-CPC localized both to overlap regions and to single microtubule segments within bundles. As sliding progressively narrowed the overlap width, CPC molecules were retained within overlap regions, while additional CPC molecules were actively transported along single microtubules into the shrinking overlaps (Figure S3C, Movie 1A). This resulted in depletion of CPC from single microtubules and progressive concentration within overlap regions. However, some CPC molecules remained on single microtubules at steady state. In contrast, when both KIF4A and KIF20A were present, CPC transport to overlap regions was more efficient, leading to more complete accumulation at overlaps, in addition to retention of CPC at overlaps during sliding (Figure S3C, Movie 1B). As expected, in the absence of motor proteins, CPC showed no preferential accumulation at overlaps, exhibiting uniform distribution along overlaps and single microtubules over time (Figures S3C, Movie 1C). When KIF20A was present as the sole motor, sliding was minimal. While a low level of CPC transport is observed, it was insufficient to promote accumulation at overlaps (Figures S3C, Movie 1D). This finding is unexpected given that the CPC-KIF20A module has long been proposed as a primary mechanism for CPC midzone localization, and PRC1 is proposed to serve as a recruitment scaffold for several midzone proteins.

Together, these systematic bottom-up reconstitutions identify human KIF4A as a key midzone factor that promotes CPC enrichment at PRC1-crosslinked antiparallel overlaps. Moreover, KIF4A and KIF20A act synergistically to maximize CPC accumulation *in vitro*. Thus, PRC1, KIF4A, KIF20A, and CPC together define the minimal module required for CPC enrichment at antiparallel microtubule overlaps.

### KIF4A is required for CPC localization at the spindle midzone in anaphase cells

We investigated the role of KIF4A in CPC enrichment at the spindle midzone during anaphase. To this end, we generated a CRISPR-edited DLD1 cell line in which endogenous KIF4A was tagged with an auxin-inducible degron (IAA) for conditional degradation and a SNAP tag for in-cell visualization. Anaphase enrichment was achieved using a previously established synchronization protocol^34^. Briefly, cells were arrested at the G1/S transition by thymidine treatment for 24 hours, released, and then incubated for 16 hours with the Cdk1 inhibitor RO3306 to synchronize cells in G2. Following release from G2 arrest, 5-Ph-IAA was added for 1 hour to deplete KIF4A prior to fixation (Figure S4A). Western blot analysis confirmed that auxin treatment reduced endogenous KIF4A levels by 87% within 1 hour (Figure 4A). Fixed cells were stained for endogenous PRC1 to visualize midzone microtubules, and either Aurora B or INCENP as CPC markers.

**Figure 4:**
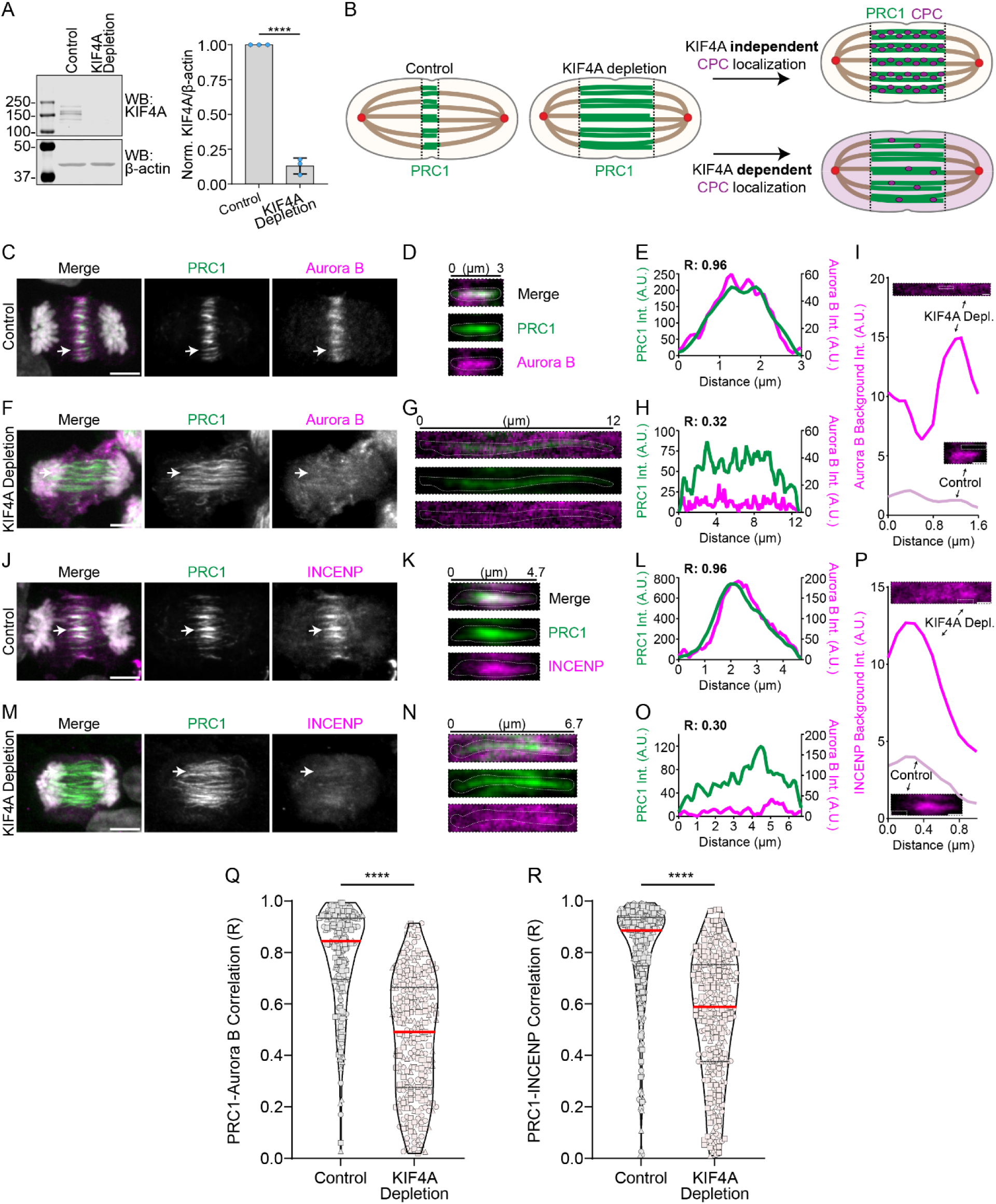
KIF4A depletion disrupts CPC enrichment at PRC1-crosslinked bundles at anaphase. (A) Western blot of endogenous KIF4A-SNAP in control and KIF4A-depleted cells. β-actin was used as a loading control. The bar graph shows the normalized KIF4A-to-β-actin intensity in control and KIF4A-depleted cells (N=3). (B) Schematic of potential outcomes for CPC localization at PRC1-bundles in anaphase cells upon KIF4A depletion. (C) Representative immunofluorescence maximum intensity projection image of endogenous PRC1 (green) and Aurora B (magenta) in control cells. Arrows indicate the individual PRC1 bundle shown in D. The scale bar is 5 μm. (D-E) Zoomed in view (D) and intensity profile (E) of an individual PRC1 bundle (green) and the associated Aurora B (magenta) signal from C. Correlation coefficient (R) is indicated. The line shown in D was used to generate the intensity profile in E. (F) Representative immunofluorescence maximum intensity projection image of endogenous PRC1 (green) and Aurora B (magenta) in KIF4A-depleted cells. Arrows indicate the individual PRC1 bundle shown in G. The scale bar is 5 μm. (G-H) Zoomed in view (G) and intensity profile (H) of an individual PRC1 bundle (green) and the associated Aurora B (magenta) signal from F. Correlation coefficient (R) is indicated. The line shown in G was used to generate the intensity profile in H. (I) The intensity profile shows the background intensity of Aurora B (magenta) in control (C) and KIF4A-depletion (F). The white boxes indicate the positions of the lines used to plot the intensity profiles for Aurora B. The scale bars are 1 μm. (J) Representative immunofluorescence maximum intensity projection image of endogenous PRC1 (green) and INCENP (magenta) in control cells. Arrows indicate the individual PRC1 bundle shown in K. The scale bar is 5 μm. (K-L) Zoomed in view (K) and intensity profile (L) of an individual PRC1 bundle (green) and the associated INCENP (magenta) signal from J. Correlation coefficient (R) is indicated. The line shown in K was used to generate the intensity profile in L. (M) Representative immunofluorescence maximum intensity projection image of endogenous PRC1 (green) and INCENP (magenta) in KIF4A-depleted cells. Arrows indicate the individual PRC1 bundle shown in N. The scale bar is 5 μm. (N-O) Zoomed in view (N) and intensity profile (O) of an individual PRC1 bundle (green) and the associated INCENP (magenta) signal from M. Correlation coefficient (R) is indicated. The line shown in N was used to generate the intensity profile in O. (P) The intensity profile shows the background intensity of INCENP (magenta) in control (J) and KIF4A-depletion (M). The white boxes indicate the positions of the lines used to plot the intensity profiles for INCENP. The scale bars are 1 μm. (Q) Violin plot shows the correlation (R) between PRC1 and Aurora B signal in control and KIF4A-depleted anaphase cells. Horizontal lines indicate the median (red line) and 25^th^ and 75^th^ percentiles (grey dotted lines). Control: N=234 (median [IQR]: 0.84 [0.69,0.93]), and KIF4A-depleted cells: N=267 (median [IQR]: 0.49 [0.27,0.66]). ****p < 0.0001. (R) Violin plot shows the correlation (R) between PRC1 and INCENP signal in control and KIF4A-depleted anaphase cells. Horizontal lines indicate the median (red line) and 25^th^ and 75^th^ percentiles (grey dotted lines). Control: N=359 (median [IQR]: 0.88 [0.74,0.93]), and KIF4A-depleted cells: N=299 (median [IQR]: 0.58 [0.37,0.75]). ****p < 0.0001.

A key experimental challenge in studying the function of KIF4A in CPC localization is that KIF4A also regulates tubulin polymerization dynamics, and its depletion leads to an elongated spindle midzone (Figure 4B)^37,38,42^. However, since KIF4A functions downstream of PRC1, PRC1-crosslinked midzone microtubule bundles still form in the absence of KIF4A (Figure 4B)^37,38^. In addition, midzone localization of other cytokinetic factors (e.g., MKLP1, CENPE) closely parallels PRC1 localization^37^. Based on these observations, we reasoned that if CPC recruitment is independent of KIF4A, its localization pattern would mirror that of PRC1 on the extended midzone bundles (Figure 4B).

In control anaphase cells, PRC1 and Aurora B were highly enriched and colocalized at the spindle midzone, consistent with prior reports (Figure 4C-D)^33,34,37^. KIF4A-depleted cells displayed an elongated spindle midzone with wider PRC1 bundles, consistent with published reports^37,38,42^. We found that Aurora B failed to properly colocalize with PRC1 under these conditions, appearing predominantly cytoplasmic and greatly reduced from PRC1-marked antiparallel bundles (Figure 4F-G).

To quantify Aurora B localization on PRC1 bundles, we performed line scan analysis of individual midzone bundles in both control and KIF4A-depleted cells and quantified the fluorescence signal for PRC1 and Aurora B in each bundle (Figure 4D-E, 4G-H). Correlation analysis revealed strong colocalization of Aurora B with PRC1 in control cells (R = 0.78, IQR: 0.69,0.93; Figure 4Q). In KIF4A-depleted cells, this correlation was reduced by ∼40% (R = 0.49, IQR: 0.27,0.66; Figure 4Q). Importantly, the observed mislocalization was not a consequence of lower CPC density on elongated bundles but reflected a failure of CPC recruitment to PRC1 bundles in the absence of KIF4A, as evidenced by the increased cytoplasmic signal of Aurora B (Figure 4I). The total Aurora B levels remained unchanged following KIF4A depletion (Figure S4B-C), indicating that the reduction in midzone localization was not due to altered protein levels.

To further validate these findings, we analyzed INCENP localization as an independent CPC marker. In control cells, INCENP strongly colocalized with PRC1 at the midzone (R = 0.88, IQR 0.74,0.93; Figure 4J-L, 4R), whereas in KIF4A-depleted cells, INCENP accumulated in the cytoplasm with diminished association with PRC1 bundles (Figures 4M-O). The correlation between INCENP and PRC1 was reduced by 34% (R = 0.558, IQR: 0.37,0.75); Figure 4R). As observed with Aurora B, the cytoplasmic levels of INCENP increased in the KIF4A-depleted conditions (Figure 4P). These findings demonstrate that depletion of KIF4A significantly impacts CPC recruitment to the PRC1-crosslinked midzone microtubule bundles.

Collectively, in agreement with our *in vitro* reconstitution results, these data establish KIF4A as a key midzone protein governing CPC localization. Since KIF20A is required for the dissociation of CPC from chromosomes and its microtubule binding at anaphase onset^30–34^, these findings suggest that KIF4A and KIF20A act collaboratively to ensure CPC midzone localization at anaphase.

## DISCUSSION

### Reconstitution of a minimal CPC midzone enrichment module

Central to this study was the purification of the human CPC (Figure 1B–D). Our systematic bottom-up reconstitutions reveal the mechanism by which CPC selectively accumulates at antiparallel microtubule overlaps. The findings rule out several prevailing models. First, we found that CPC alone does not enrich significantly at microtubule bundles (Figure 1H-J). Second, we tested a model in which KIF20A’s microtubule bundling activity (Figure 2F-H) and its interaction with CPC underlie its transport and enrichment at microtubule overlaps. However, we found that the CPC-KIF20A module does not enrich CPC at overlaps (Figures 2F-H, 3D, S3C). Finally, the reconstitutions showed that PRC1, which crosslinks antiparallel microtubules and is highly enriched at the overlap, cannot directly recruit CPC (Figures 2J-L, 3E, S3C). We identified KIF4A as the missing link: it binds both PRC1 and CPC, acting as a molecular link between CPC and PRC1-crosslinked antiparallel microtubule overlaps (Figures 3B, S3C, 3G, 5A).

The inclusion of KIF20A markedly enhances CPC accumulation by KIF4A at PRC1-crosslinked microtubule overlaps (Figures 3C, 3G, S3C). The data suggest a mechanism where KIF20A, which binds CPC with high affinity^30–34^, efficiently transports CPC along single microtubules flanking the overlap region. Once delivered, CPC is retained by KIF4A. This is because KIF4A is concentrated at the overlaps through high-affinity binding with PRC1, which, in turn, selectively accumulates at antiparallel overlaps. Thus, the division of labor between KIF20A and KIF4A underlies the selective enrichment of CPC at antiparallel overlaps. Importantly, while each component in this network can crosslink microtubules, a sequential set of interactions is required to achieve robust and spatially precise CPC accumulation at antiparallel overlaps (Figure 5A).

**Figure 5:**
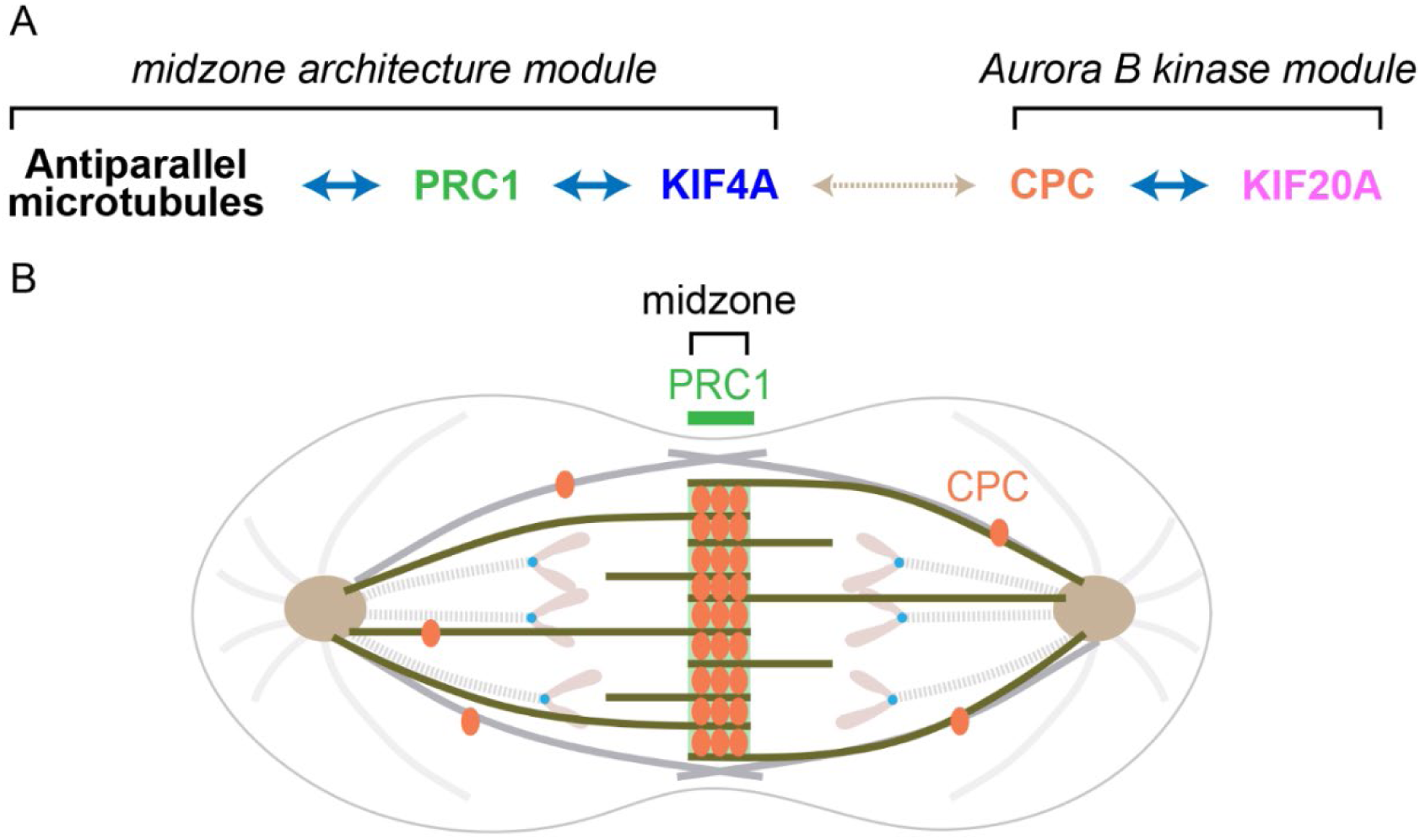
The collective activity of KIF20A and KIF4A controls CPC localization at antiparallel midzone microtubules. (A) A “minimal CPC midzone recruitment module” ensures the localization of CPC at the spindle midzone through a linear interaction network that links the antiparallel microtubules to the CPC. The link between the midzone architecture module and the CPC-KIF20A module is formed by a direct interaction between KIF4A and CPC. (B) In the context of the spindle midzone, KIF20A transports the CPC outside the midzone, while KIF4A would retain and concentrate CPC within the sliding PRC1-anti-parallel microtubules of the midzone (green). In the schematic, CPC is shown as orange ovals. The heterogeneity of the microtubule network is illustrated using different colors for distinct microtubule subsets. Microtubules that form the midzone: greenish brown; microtubules forming the cortical array: gray; astral microtubules: light gray; kinetochore microtubules: dotted light gray. PRC1 (green) marks the spindle midzone.

### Collaborative role of KIF20A and KIF4A for midzone localization of CPC in anaphase cells

Prior studies have shown that KIF20A is required for the dissociation of CPC from chromosomes at anaphase onset in mammalian cells^30–34^. Replacing wild-type KIF20A with a rigor mutant results in CPC localization predominantly on interpolar microtubules between the chromosomes and the edge of the midzone, with reduced levels at the central overlap region in anaphase^30,31,34^. These findings suggest that KIF20A motility or turnover on interpolar microtubules outside the midzone plays an important role in CPC delivery to the midzone. Results from our conditional depletion studies of KIF4A show that in the absence of KIF4A, KIF20A-driven transport alone cannot localize CPC to the extended midzone at anaphase; instead, CPC remains largely cytoplasmic (Figure 4). These data suggest that the concentrated pool of KIF4A at the midzone acts as the site-specific scaffold for CPC recruitment and retention. Thus, CPC phosphorylates KIF4A to activate it for its function as a midzone length regulator^5,39^, and the CPC-KIF4A interaction concentrates CPC as the midzone width narrows during microtubule sliding in anaphase. Collectively, these data are consistent with a mechanism where KIF20A activity is important to deliver CPC to the midzone and KIF4A is required for CPC retention at PRC1-crosslinked midzone microtubules in anaphase cells (Figure 5A-B).

### Strategies for effective CPC navigation within diverse spindle architectures

Midzone localization of CPC is a conserved step in mitosis. Since spindle size and architecture vary across organisms and cell types, distinct solutions are likely to have evolved to achieve CPC enrichment at the midzone in different microtubule networks. Intriguingly, multimotor transport systems may be a general theme. In *Xenopus* egg extracts, the KIF4A homolog and an embryonic KIF20A paralog (KIF20AE) are both required for CPC localization to the aster-aster interaction zone (AAIZ) of large asters. In this system, KIF20AE is critical for CPC recruitment to asters, and the KIF4A homolog is important for CPC transport along astral microtubules towards the AAIZ^40^. In budding yeast, kinesins Kip1 and Kip3 associate with CPC and are proposed to capture diffusing complexes at the midzone^44^. In *Trypanosoma brucei*, Aurora B and INCENP are constitutively bound to KIN-A and KIN-B, where KIN-A’s ATPase activity is essential for midzone localization and KIN-B lacks the features of a functional kinesin motor^45^. Further reconstitutions will reveal how the properties of these motors are tuned to ensure CPC localization at antiparallel microtubule overlaps within the different spindle contexts.

## MATERIAL AND METHODS

### Plasmids

The cDNA sequences encoding for human full-length INCENP (Inner Centromere Protein) and Aurora B kinase were codon optimized and subcloned into a pFastBac dual expression vector (ThermoFisher Scientific, 10712024) using the In-Fusion HD Cloning kit (Takara Bio, 638910). INCENP protein was followed by a PreScission Protease cleavable C-terminal Flag-tag. To insert the INCENP-PreScission-Flag sequence, we used the following primers: 5’-GGTCCGAAGCGCGCGGAATT-3’ and 5’-GGATCCGCGCCCGATGGTG-3’ were used to amplify the plasmid and 5’-ATCGGGCGCGGATCCATGGGCACCACCGCTCCCG-3’ and 5’-CCGCGCGCTTCGGACCTTACTTGTCATCGTCGTCCTTGTAGTCGGGCCCCTGG-3’ to amplify and insert the sequence for INCENP-PreScission-Flag. The new construct was then used as a template to insert the sequence of Aurora B tagged with a 6xHis tag, GFP and a tobacco etch virus (TEV) cleavable site at its N-terminus. Primers 5’-GTGATCAAGTCTTCGTCGAGTGAT-3’ and 5’-GAGATGGGGGAGGCTAACTGAAA-3’ were used to amplify the new construct, and primers 5’-AGCCTCCCCCATCTCTTAAGCCACGGATTGCAGAG-3’ and 5’-CGAAGACTTGATCACATGTCGTACTACCATCACCATCACCATCACGA-3’ were used to amplify and insert the sequence of 6xHis-GFP-Aurora B. When needed, the sequence of GFP was deleted using the primers: 5’-GGCGCCCTGAAAATACAGGTTTTCGGTCGTTGGGA-3’ and 5’-GGCGCCCTGAAAATACAGGTTTTCGGTCGTTGGGA-3’. Similarly, the sequences encoding for human Borealin and Survivin were subcloned into a pFastBac dual expression vector using the In-Fusion HD Cloning kit. Unlabeled or Halo-tagged Borealin protein was followed by a tobacco etch virus (TEV) cleavable C-terminal Twin-Strep-Tag. The Halo tag was inserted between Borealin and TEV site. To generate the plasmid, we used the following primers: 5’-ACGACCGAAAACCTGTATTTTCAGG-3’ and 5’-CTTGTGGGTACGGATGGAGG-3’ were used to amplify the plasmid, and 5’-ATCCGTACCCACAAGGAAATCGGTACTGGCTTTCC-3’ and 5’-CAGGTTTTCGGTCGTACCGGAAATCTCCAGAGTAGACAG-3’ to amplify and insert the sequence for the Halo tag. Survivin protein was untagged.

KIF20A was cloned into a modified pFastBac expression vector, incorporating a 6xHis and Sumo tag upstream of the protein, with or without a downstream GFP tag. A TEV cleavage site was positioned between the Sumo tag and KIF20A. For the unlabeled KIF20A: the sequence for KIF20A was amplified from clone ID 4538604 (in pOTB7 vector) from Open Biosystems using the primers 5’-TCAGGGCGGCTCTAGTATGTCGCAAGGGATCCTTTCTCC-3’ and 5’-CCGCGGCCGCTCTAGTTAGTACTTTTTGCCAAAAGGCCC-3’. Both the amplified PCR product and expression vector were digested with XbaI and subsequently ligated with the In-Fusion HD Cloning kit. For the KIF20A tagged with GFP at the C-terminus: as above, the sequence for KIF20A was amplified using the primers 5’-TCAGGGCGGCTCTAGTATGTCGCAAGGGATCCTTTCTCC-3’ and 5’-CCTTGCTCACCATAGTGTACTTTTTGCCAAAAGGCCC-3’. The amplified PCR product and expression vector with GFP were digested with XbaI and KpnI. The final construct was generated using the In-Fusion HD Cloning kit.

### Protein expression and purification

For CPC expression, plasmids encoding for INCENP/Aurora B and Borealin/Survivin were co-expressed in Sf9 insect cells using the Bac-to-Bac Baculovirus expression system. For CPC-GFP, Halo-CPC-GFP, and Halo-CPC, cells were grown in HyClone CCM3 (Cytiva, SH30065.02) and expressed for 68-70 h at 27 °C. Before pelleting the cells at 4,000 x g for 20 min at 4°C, they were incubated with 80 μM ZnCl_2_ for 16-18 h. Cell pellets were resuspended in lysis buffers (50 mM Potassium Phosphate pH 8.0, 500 mM NaCl, 5% (v/v) glycerol, 100 μM ATP, 30 mM Imidazole, 2 mM tris(2-carboxyethyl)phosphin (TCEP, Thermo Fisher Scientific, 77720), 1 mM phenylmethanesulfonyl fluoride (PMSF), 0.15% (v/v) Tween-20, 0.5% (v/v) Igepal 630, 75U of benzonase (Sigma-Aldrich, 70746-3), 0.1 mg/ml lysozyme and 1x HALT protease inhibitor cocktail (Thermo Fisher Scientific, PI78439), lysed by sonication and clarified by centrifugation at 61,740 x g (Ti-70.1 rotor) for 40 min at 4°C. Supernatants were incubated with Ni-NTA resin (Qiagen, 30230) for 2 h at 4°C. Bound protein complexes were washed with wash buffer (50 mM Potassium Phosphate pH 8.0, 1 M NaCl, 5% (v/v) glycerol, 100 μM ATP, 30 mM Imidazole, 0.5 mM tris(2-carboxyethyl)phosphin (TCEP), 0.15% (v/v) Tween-20 and eluted with 50 mM Potassium Phosphate pH 8.0, 500 mM NaCl, 5% (v/v) glycerol, 100 μM ATP, 250 mM Imidazole, 0.5 mM tris(2-carboxyethyl)phosphin (TCEP), 0.15% (v/v) Tween-20. Eluted protein was further purified by size exclusion chromatography (SEC) using Superose 6 Increase 10/300 GL column (Cytiva, 29-0915-96) in buffer containing 50 mM Potassium Phosphate pH 8.0, 500 mM NaCl, 10% (v/v) glycerol, 100 μM ATP, 10 mM 2-Mercaptoethanol, 4 mM DTT, and 0.15% (v/v) Tween-20. Peak fractions were incubated with PreScission protease overnight at 4°C followed by incubation with Strep beads (Strep-Tactin Sepharose, IBA LifeSciences, 2-1201-010 and 2-1250-010) for 2-3 h at 4°C. Bound protein was eluted in buffer containing 50 mM Potassium Phosphate pH 8.0, 500 mM NaCl, 10% (v/v) glycerol, 100 μM ATP, 10 mM 2-Mercaptoethanol, 4 mM DTT, 0.15% (v/v) Tween-20 supplemented with 10 mM d-Desthiobiotin (Sigma-Aldrich, D1411). Eluted proteins were incubated with TEV protease overnight at 4°C. When applicable, Halo-CPC was labeled with Janelia Fluor 646 HaloTag Ligand (Promega, GA1120) by incubating the protein with the ligand overnight at 4°C (1:6 molar ratio, respectively). Cleaved proteins were further purified by size exclusion chromatography (SEC) using Superose 6 Increase 10/300 GL column in buffer containing 50 mM Potassium Phosphate pH 8.0, 500 mM NaCl, 10% (v/v) glycerol, 100 μM ATP, 10 mM 2-Mercaptoethanol, 4 mM DTT and 0.15% (v/v) Tween-20. Peak fractions were concentrated, supplemented with 30% sucrose, and proteins flash-frozen in liquid nitrogen and stored at -80°C (Figures 1B, S1).

The presence of all protein components in CPC was verified by Western blots. CPC complexes were loaded into 4-20% SDS-PAGE gels and transferred to a nitrocellulose membrane. Subsequently, membranes were blocked for 1 h at room temperature with a blocking buffer (5% w/v nonfat dry milk and 1% w/v BSA in phosphate-buffered saline (PBS) 1x pH 7.4). Membranes were washed with PBS/T (PBS 1x pH 7.4/0.1% Tween-20) and incubated overnight at 4°C in 2% (w/v) BSA with PBS/T with a mouse antibody against INCENP (Thermo Fisher Scientific, 39-2800), rabbit antibody against Aurora B (Abcam, ab2254), mouse antibody against Borealin (Santa Cruz Biotechnology, sc-376635) and mouse antibody against Survivin (Santa Cruz Biotechnology, sc-17779). All antibodies were diluted 1:5000. Subsequently, the membranes were washed with PBS/T and incubated with goat anti-rabbit IRDye 800CW (LICORbio, 926-32211) and goat anti-mouse IRDye 680LT (LICORbio, 926-68020) secondary antibodies for 1 h at room temperature. Subsequently, membranes were washed with PBS/T and imaged with an Odyssey CLx imaging system (LI-COR).

For KIF20A or KIF20A-GFP expression, plasmids encoding KIF20A or KIF20A-GFP were co-expressed in Sf9 insect cells using the Bac-to-Bac Baculovirus expression system. Insect cells were grown in HyClone CCM3 and expressed proteins for 68-70 h at 27 °C. After the expression, cells were centrifuged at 4,000 x g for 20 min at 4°C. Cell pellets were resuspended in lysis buffers (50 mM Potassium Phosphate pH 8.0, 500 mM NaCl, 10% (v/v) glycerol, 1 mM MgCl_2_, 100 μM ATP, 30 mM Imidazole, 2 mM tris(2-carboxyethyl)phosphin (TCEP), 1 mM phenylmethanesulfonyl fluoride (PMSF), 0.15% (v/v) Tween-20, 0.5% (v/v) Igepal 630, 75U of benzonase, 0.2 mg/ml lysozyme, 1x HALT protease inhibitor cocktail, and 2 mM Benzamidine Hydrochlorode, lysed by sonication and clarified by centrifugation at 61,740 x g (Ti-70.1 roror) for 40 min at 4°C. Subsequently, supernatants were incubated with Ni-NTA resin for 1 h and 30 min at 4°C. Bound protein complexes were washed with wash buffer (50 mM Potassium Phosphate pH 8.0, 500 mM NaCl, 10% (v/v) glycerol, 1 mM MgCl_2_, 100 μM ATP, 30 mM Imidazole, 2 mM tris(2-carboxyethyl)phosphin (TCEP), and 0.15% (v/v) Tween-20. Bound proteins were then eluted with 50 mM Potassium Phosphate pH 8.0, 500 mM NaCl, 10% (v/v) glycerol, 1 mM MgCl_2_, 100 μM ATP, 400 mM Imidazole, 0.5 mM tris(2-carboxyethyl)phosphin (TCEP), and 1x HALT protease inhibitor cocktail. Eluted proteins were concentrated, supplemented with 2 mM Ethylenediaminetetraacetic acid (EDTA) and 5 mM 2-Mercaptoethanol and incubated with TEV protease overnight at 4°C. Cleaved proteins were further purified by size exclusion chromatography (SEC) using Superose 6 Increase 10/300 GL column in buffer containing 50 mM Potassium Phosphate pH 8.0, 500 mM NaCl, 10% (v/v) glycerol, 1 mM MgCl_2_, 200 μM ATP, 5 mM 2-Mercaptoethanol. Peak fractions were concentrated, supplemented with 30% sucrose, and proteins flash-frozen in liquid nitrogen and stored at -80°C (Figures 2, 3, S2).

For the expression and purification of GFP-PRC1 isoform 2 and KIF4A, we followed the protocols previously described in Subramanian et al, 2010^41^ and 2013^43^, respectively (Figures 2, 3, S2, S3).

All concentrations refer to monomeric proteins.

### Mass Photometry assay

To determine the molecular weight of the purified CPC-GFP, we utilized mass photometry. Refeyn Two MP Mass Photometer (Refeyn) was calibrated with calibration markers, and subsequently, we determined its molecular weight at a final concentration of 10 nM (Figure 1C).

### Pull-down assays

To test for the CPC-KIF20A interaction, we performed GFP-pull down assays (Figure S2C). First, we incubated CPC-GFP (2.5 μg) with GFP-trap agarose beads (ChromoTek, gta-10) pre-equilibrated with pull-down buffer (BRB80 pH 6.8 (80 mM Pipes-KOH pH 6.8, 2 mM MgCl_2_, 1 mM EGTA, 50 mM KCl, 100 μM ATP, 0.1% (w/v) Triton X-100, 1x HALT protease inhibitor cocktail, 1 mM TCEP and 5% (w/v) sucrose for 1 h at 4°C. Subsequently, beads were washed twice with pull-down buffer and incubated for 2 h at 4°C with unlabeled KIF20A (2.5 μg). Following incubation, bound protein complexes were washed five times with pull-down buffer before eluting with SDS loading buffer and boiled. Subsequently, equal amounts of eluted protein complexes were loaded into 12.5% SDS-PAGE gels and stained with Coomassie R-250 or transferred to a nitrocellulose membrane. Membranes were blocked for 1 h at room temperature with a blocking buffer (5% nonfat dry milk and 1% BSA in phosphate-buffered saline (PBS) 1x pH 7.4). Then, they washed with PBS/T (PBS 1x pH 7.4/0.1% Tween-20) and incubated for 1 h at room temperature in 2% (w/v) BSA with PBS/T with primary antibodies: mouse anti-GFP at 1:5000 dilution (Santa Cruz Biotechnology, sc-9996) and mouse anti-Survivin at 1:2000 dilution. Subsequently, the membranes were washed with PBS/T and incubated with goat anti-rabbit IRDye 800C and goat anti-mouse IRDye 680LT secondary antibodies for 1 h at room temperature. Finally, the membranes were washed with PBS/T and imaged with an Odyssey CLx imaging system (LI-COR). After imaging, the membranes were incubated with 0.2N NaOH for 10 min at room temperature to remove the antibodies and washed three times with MilliQ water. Then membranes were blocked and incubated overnight with primary antibodies: mouse anti-Flag at 1:2000 dilution (Sigma-Aldrich, F3165) and rabbit anti-Strep at 1:2000 (GenScrip, A00626). Membranes were washed and incubated with goat anti-mouse IRDye 680LT secondary antibodies for 1 h at room temperature, before imaging.

### Microtubule polymerization

Taxol-stabilized and biotinylated (9%, Cytoskeleton, T333P) GMPCPP microtubules labeled with HiLyte 488 (9%, Cytoskeleton, TL488M), X-rhodamine (9%, Cytoskeleton, TL620M), or HiLyte 647 (9%, Cytoskeleton, TL670M) and X-rhodamine-labeled (9% or 20%), non-biotinylated taxol-stabilized GMPCPP microtubules were prepared as previously described^14,41,43^. Briefly, the GMPCPP (Jena Bioscience, NU-405) microtubule seeds were incubated for 1 h and 30 min at 37°C. Subsequently, polymerized microtubules were mixed with warm BRB80 pH 6.8 (80 mM Pipes-KOH pH 6.8, 2 mM MgCl_2_, 1 mM EGTA) and centrifuged at 244,900 x g for 20 min at 30°C. Pelleted microtubules were resuspended in BRB80 pH 6.8 supplemented with 20 μM taxol (Selleck Chemicals, S1150) and centrifuged as above. Finally, pelleted microtubules were resuspended in BRB80 pH 6.8 supplemented with 20 μM taxol and stored at room temperature in the dark.

### TIRF microscopy assays

The imaging chambers for the TIRF assays were prepared and assembled as before^14,41,43,46^. Briefly, microscopy glass slides (Electron Microscopy Sciences, 63793-01) and coverslips (Electron Microscopy Sciences, 63787-01) were cleaned, functionalized, and treated with biotinylated PEG (Laysan Bio, Biotin-PEG-SVA-5000) and non-biotinylated PEG (Laysan Bio, mPEG-SVA-5000), respectively. We then used double-sided adhesive sheets (Soles2dance, 9474-08×12 3M 9474LE 300LSE) and a paper cutter (Silhouette, Portrait 3) to prepare custom imaging chambers with a volume of approximately 10 μl.

*TIRF assays for microtubule overlaps:* To make microtubule bundles with CPC, KIF20A, or PRC1 (Figures 1, 2, 3, S1F-G, S3C), we used the protocol described in Subramanian et al, 2010^41^. First, we treated the imaging chambers with neutravidin (0.2 mg/ml, Invitrogen, A2666) for 5-10 min. Subsequently, the imaging chambers were washed with assay buffer (BRB80 pH 6.8, 5 mM TCEP, 1 mM DTT, 5% (w/v) sucrose and incubated with κ-casein (0.5 mg/ml, Sigma-Aldrich, C0406) in assay buffer supplemented with 20 μM taxol for 5 min. Then, imaging chambers were incubated with biotinylated taxol-stabilized and fluorescent-labeled GMPCPP microtubules for 15 min. After incubation, imaging chambers were washed with κ-casein (0.5 mg/ml) in assay buffer supplemented with 20 μM taxol to remove unbound microtubules. Next, immobilized microtubules were incubated with different concentrations of Halo 646-CPC or unlabeled KIF20A or 1 nM GFP-PRC1 isoform 2 for 5 min. Subsequently, unbound proteins were removed by washing the imaging chambers, which were then incubated with X-rhodamine-labeled, non-biotinylated taxol-stabilized GMPCPP microtubules for 7 min, to allow for microtubule overlap formation. Finally, the imaging chambers were washed twice with κ-casein (0.5 mg/ml) in assay buffer supplemented with 20 μM taxol, and the reaction was initiated by flowing into the chambers: i) different concentrations of Halo 646-CPC (Figures 1G-H, S1F-G), (ii) different concentrations of Halo 646-CPC and unlabeled KIF20A (Figure 2E-F), (iii) different concentrations of Halo 646-CPC with 1 nM GFP-PRC1 isoform 2 (Figure 2I-J) and (iv) 30 pM or 2.5 nM Halo 646-CPC with 15 nM unlabeled KIF20A or 15 nM KIF4A or both motors and 1 nM GFP-PRC1 isoform 2 (Figures 3A-E, S3C) in assay buffer supplemented with 20 μM taxol, 50 mM KCl, 0.5 mg/ml κ-casein, 1 mM ATP, 1 mM 6-Hydroxy-2,5,7,8-tetramethylchromane-2-carboxylic acid (Trolox, Sigma-Aldrich, 238813), 0.4 mg/ml glucose oxidase (Sigma-Aldrich, G2133), 0.2 mg/ml catalase (Sigma-Aldrich, C1345), 20 mM D-glucose (Sigma-Aldrich, G8270) and 71.5 mM 2-Mercaptoethanol. For making microtubule overlaps with KIF20A-GFP (Figure 2A-B), we followed the same protocol but the final reaction was consisted of different concentrations of KIF20A-GFP in assay buffer supplemented with 20 μM taxol, 50 mM KCl, 0.5 mg/ml κ-casein, 100 μM ATP, 1 mM Trolox, 0.4 mg/ml glucose oxidase, 0.2 mg/ml catalase, 20 mM glucose and 71.5 mM 2-Mercaptoethanol. Imaging chambers were sealed, and still images were acquired after 10 min of incubation from multiple imaging fields (Figures 1, 2) or real-time imaging was performed immediately for 20 minutes at 5 seconds/frame (Figures 3, S3C, and Movie 1). In this case, we collected images from multiple imaging fields for quantification immediately upon completion of the experiment.

*TIRF assays for microtubule binding assays:* To determine the microtubule binding affinity of Halo-CPC-GFP, we incubated the imaging chambers with neutravidin (0.2 mg/ml) for 10 min. After washing with assay buffer supplemented with 20 μM taxol, imaging chambers were incubated with κ-casein (0.5 mg/ml) in assay buffer supplemented with 20 μM taxol for 2 min. Taxol-stabilized GMPCPP X-Rhodamine biotinylated microtubules were then flown into the imaging chambers for 15 min. After the incubation, the imaging chambers were washed with assay buffer supplemented with 20 μM taxol, and incubated with κ-casein (0.5 mg/ml) in assay buffer supplemented with 20 μM taxol for 2 min. Reactions were initiated by flowing into the chambers different concentrations of Halo-CPC-GFP in assay buffer supplemented with 20 μM taxol, 50 mM KCl, 0.5 mg/ml κ-casein, 1 mM ATP, 0.4 mg/ml glucose oxidase, 0.2 mg/ml catalase, 20 mM glucose, and 71.5 mM 2-Mercaptoethanol. Imaging chambers were sealed, and still images were acquired after 10 min of incubation from multiple imaging fields (Figure 1E).

TIRF assays for CPC’s microtubule binding were performed with the Nikon Ti-E inverted microscope, equipped with a Ti-ND6-PFS perfect focus system, an APO TIRF 100x oil/1.49 DIC objective (Nikon), a Nikon-encoded x-y motorized stage and a piezo z-stage, and an EMCCD camera (Andor iXon Ultra, DU-897U-CSO-#BV). TIRF assays for CPC’s localization at microtubule overlaps were performed on a customized inverted Nikon Eclipse Ti microscope equipped with a two-stepping motor actuators (SGSP-25ACTR-B0; Sigma Koki) mounted on a KS stage (model KS-N), a 100× (1.45 N.A.) oil objective (Plan Apo ƛ; Nikon), a reflection-based autofocus unit (Focustat4), and Andor iXon EM cameras, DU-897E. μManager 2.0 was used for acquisition^47^.

### Cell culture and Immunofluorescence microscopy

To generate the KIF4A knock-in cell line, an sgRNA targeting Kif4A (5’-CACCGCATCTCCCAAGCCAGACTG-3’ and 5’-AAACCAGTCTGGCTTGGGAGATGC-3’) was cloned into the pX330 vector and sequence-verified (pMB1345). For the homology repair (HR) construct, each homology region (HR) was amplified separately (3HA 5’-AGTTGGAGTCATCATCTCTACCACCAGT-3’ and 5’-TCCTAGGGTAGCAGAGCAACATAAAGACACA-3’, and for 5HR: 5’-TGACCTCCCCAAGCAGGGCC-3’ and 5’-GTGGGCCTCTTCTTCGATAGGGGAGC-3’). A selection cassette encoding IAA17-SNAP-P2A-Neo was amplified from pMB1320. The three PCR products (two HR arms and the selection cassette) were combined and assembled using NEB HiFi DNA Assembly (New England Biolabs, E2621). The assembled product was then amplified using the outermost primers, and the resulting PCR product was used directly for transfection. DLD1 cells were transfected with 1.5 µg of pMB1345 and 1.5 µg of the HR PCR product using Lipofectamine 3000 (Thermo Fisher Scientific, L3000015). Three days post-transfection, cells were selected in neomycin-containing medium for two weeks. Single-cell colonies were isolated and screened by genomic DNA PCR to identify positive clones. To introduce Tir1F74G into these cells, the Tir1F74G HR construct (pMB1398) was knocked into the AAVS1 locus of homozygous DLD1 KIF4A-AID-SNAP cells using the sgRNA plasmid (pMB1422). Single-cell clones were isolated, screened for successful knock-in by genomic DNA PCR, and then validated for KIF4A degradation by microscopy and Western blotting following auxin treatment.

CRISPR-edited DLD1 cells with endogenous KIF4A-AID-SNAP were cultured in RPMI-1640 Medium (Sigma-Aldrich, R8758) supplemented with 10% Fetal Bovine Serum (Sigma-Aldrich, 12306C) and 1% penicillin/streptomycin (v/v) (Gibco, Cat#15140) at 37°C, 5% CO_2_ in a humidified sterile incubator. For immunofluorescence experiments, 0.3-0.4 x 10^6^ cells were cultured in 60 mm plates with 18 mm square coverslips coated with 50 μg/ml Collagen (Advanced Biomatrix, 5005) supplemented with 2.5 mM thymidine (Sigma-Aldrich, T1895). Cells were incubated for 24 h and washed three times with RPMI-1640 Medium (supplemented with FBS and penicillin/streptomycin). Subsequently, cells were incubated with 5 μM RO3306 (Sigma-Aldrich, 217699) for 16 h to synchronize them in G2^34^. Cells were then washed three times with RPMI-1640 Medium (supplemented with FBS and penicillin/streptomycin) and further incubated with 1 μM 5-Ph-IAA (Tocris Bioscience, 7392) to induce KIF4A depletion or DMSO for control cells.

Subsequently, cells were fixed in pre-warmed at 37°C PHEM buffer (60 mM Pipes-KOH pH 6.9, 25 mM Hepes, 10 mM EDTA, and 2 mM MgCl_2_, supplemented with 4% paraformaldehyde (Thermo Fisher Scientific, 28908) and 0.2% (v/v) Triton X-100 for 10 min. Then, cells were washed with PBS pH 7.4 (Phosphate Buffered Saline). Cells were quenched with Glycine 1M pH 8.5 for 5 min and washed with PBS-T (PBS pH 7.4, 0.1% (v/v) Triton X-100). Next, cells were permeabilized in PBS-T (PBS pH 7.4, 1 % (v/v) Triton X-100) for 20 min. After the incubation, cells were washed three times with PBS-T and blocked in blocking buffer (3% w/v BSA, PBS-T) for 30 min at room temperature. Primary antibodies were diluted in blocking buffer and incubated for 2 h at room temperature. Primary antibodies used were: rabbit anti-PRC1 at 1:2000 dilution^43^, mouse anti-INCENP at 1:100 dilution, and mouse anti-Aurora B at 1:500 dilution (BD Biosciences, 611082). Subsequently, cells were washed three times with PBS-T and incubated with secondary antibodies for 1 h at room temperature. Secondary antibodies used were: Alexa Fluor 488 donkey anti-rabbit IgG at 1:1000 dilution (Thermo Fisher Scientific, A-21206) and Alexa Fluor 594 donkey anti-mouse IgG at 1:1000 dilution (Thermo Fisher Scientific, A-21203). 4’,6-Diamidino-2-Phenylindole (DAPI, Invitrogen, D3571) at 2 μg/ml was used for DNA staining. Cells were washed three times with PBS-T, followed by one wash in PBS, and finally, samples were mounted in Prolong Diamond Antifade Mountant (Molecular Probes, P36961). Z-stacks were collected every 0.4 μm step with a Nikon A1R laser-scanning confocal inverted microscope equipped with an APO TIRF 60x oil/1.49 DIC objective (Nikon) (Figure 4).

For protein quantification in cell extracts from control and KIF4A-depleted cells (Figures 4A, S4B), cells were synchronized and treated as described in the Cell culture and Immunofluorescence microscopy section. Cells were washed twice with ice-cold PBS1x and lysed in lysis buffer (50 mM Tris pH 7.4, 150 mM NaCl, 0.25% (w/v) NP-40, 0.25x Halt Phosphatase Inhibitor Cocktail (Thermo Fisher Scientific, 78420), 1x HALT protease inhibitor cocktail, 1 mM PMSF, 1 mM DTT, and 0.1% (w/v) Triton X-100. Next, the lysed cells were centrifuged at 18,200 x g for 30 min at 4°C, and the concentration of protein extracts was determined with the Bradford assay. 50 μg of protein extracts from control and KIF4A-depleted cells were loaded into 4-20% SDS-PAGE gels and transferred to a nitrocellulose membrane. As above, membranes were blocked and subsequently incubated overnight at 4°C with primary antibodies: mouse anti-KIF4 at 1:1000 dilution (Santa Cruz Biotechnology, sc-365144), mouse anti-β-actin at 1:10000 dilution (Cell Signaling Technology, 58169), mouse anti-α-tubulin at 1:20000 dilution (Sigma-Aldrich, T6199) and rabbit anti-Aurora B at 1:1000 dilution. Then, membranes were washed and further incubated with goat anti-rabbit IRDye 800CW and goat anti-mouse IRDye 680LT secondary antibodies for 1 h at room temperature. Finally, membranes were washed and imaged with an Odyssey CLx imaging system (LI-COR).

### Image and statistical analysis

All quantifications were performed with the open-source software Fiji (Fiji: an open-source platform for biological-image analysis)^48^. To quantify the fluorescence intensity of Halo-CPC-GFP on microtubules (Figure 1E-F), we manually drew segmented lines of equal width (3 pixels) along the entire length of the microtubules. Then, the average pixel intensity was calculated after subtracting background intensity using regions of interest of the same size close to each microtubule. Intensities were not analyzed for microtubules found at the edge of the camera’s field of view. Data from three technical repeats are represented as mean ± SD and fitted to a Hill equation.

To calculate the I_Overlap_/I_single_ ratio of Halo 646-CPC in Figures 1J, 2H, 2L, and 3F, and KIF20A-GFP in Figure 2D, we subtracted the background (rolling ball radius, 50 pixels), and a segmented line (3 pixels) was drawn along the entire length of the microtubule, covering both the single and overlapping regions. Then, this line was edited to define the overlap region by deleting existing points. Subsequently, both lines were converted into areas, and the “XOR” function was used to determine the region of the single microtubules. The I_Overlap_/I_single_ ratio was calculated by dividing the fluorescence intensity of the overlap region by that of the single microtubules. Statistical test for Figures 1J, 2D, 2H, 2L, and 3F: One-way ANOVA and Kruskal–Wallis H test followed by Dunn’s multiple comparison test. Intensity profiles in Figures 1, 2, 3, and S1 were generated using the plot profile function in Fiji, after background subtraction.

Kymographs in Figure S3C were generated using the KymoResliceWide plugin for ImageJ. Where it was applicable, videos were corrected for xy drift using the Fast4Dreg macro^49^. To quantify the correlation (R) between PRC1 and Aurora B or PRC1 and INCENP in Figure 4Q-R, we collected Z-stacks of control and KIF4A-depleted cells in anaphase. To ensure that all cells are in anaphase, we measured the DNA-DNA distance of individual cells between the outer edges of the DNA signal, and only cells with DNA-DNA distances of 13-17 μm were used for analysis. After background subtraction (rolling ball radius, 15 pixels), we manually traced each PRC1 bundle across the Z-stack and recorded the first and last planes where it was visible. Using the recorded planes, we created an average projection of each PRC1 bundle, where we manually drew a segmented line (5 pixels) along the entire length of the PRC1 bundle and generated intensity profiles for PRC1 and Aurora B or PRC1 and INCENP. Finally, these values were imported into GraphPad Prism, and the correlation between PRC1 and Aurora B or PRC1 and INCENP was calculated. Statistical analysis for Figure 4Q-R: Data is from three technical repeats and analyzed with unpaired t-test.

Intensity profiles in Figure 4 were generated using the plot profile function in Fiji, after background subtraction. The intensity profiles in Figures 4I and 4P were smoothed in GraphPad Prism for better visualization.

For determining the protein levels of KIF4A and Aurora B in control and KIF4A-depleted anaphase cells in Figures 4A and S4C, we measured their intensities from the western blots. To correct for differences in sample loading, the intensities of KIF4A and Aurora B were normalized against the intensities of β-actin or α-tubulin, which were used as loading controls. The corrected values were normalized to the control and plotted using GraphPad Prism. All intensities were measured using the Image Studio Lite Ver 5.2. Data is from three technical repeats and analyzed with unpaired t-test.

All the values for the N, median (IQR), mean, and S.D. for each experiment are mentioned in the figure legends. The number of repeats for each experiment is mentioned in the Materials and Methods along with the statistical tests used for analysis. All statistical analyses were performed with GraphPad Prism 10.

## ACKNOWLEDGEMENTS

This work was funded by NIH grants GM144352 (M.B.), GM122893 (M.B.), and GM155215 (R.S.). K.N. was supported by MGH ECOR Fund for Medical Discovery (FMD) Fundamental Research Fellowship Award.

## AUTHOR CONTRIBUTIONS

We thank Dr. Reito Watanabe from Dr. Michael Blower’s lab (Boston University Chobanian and Avedisian School of Medicine) for the pMB1422 and pMB1398 plasmids. K.N. and R.S. designed the research; M.B. developed the KIF4A conditional depletion cell line and advised on cell biological experiments. K.N. cloned the constructs, expressed and purified recombinant proteins, collected and analyzed TIRF and confocal microscopy data; K.N. and R.S wrote the manuscript with input from M.B.

