## Supplemental Figures_Movie caption for "A minimum module for positioning the Chromosomal Passenger Complex at the cell center for cytokinesis"

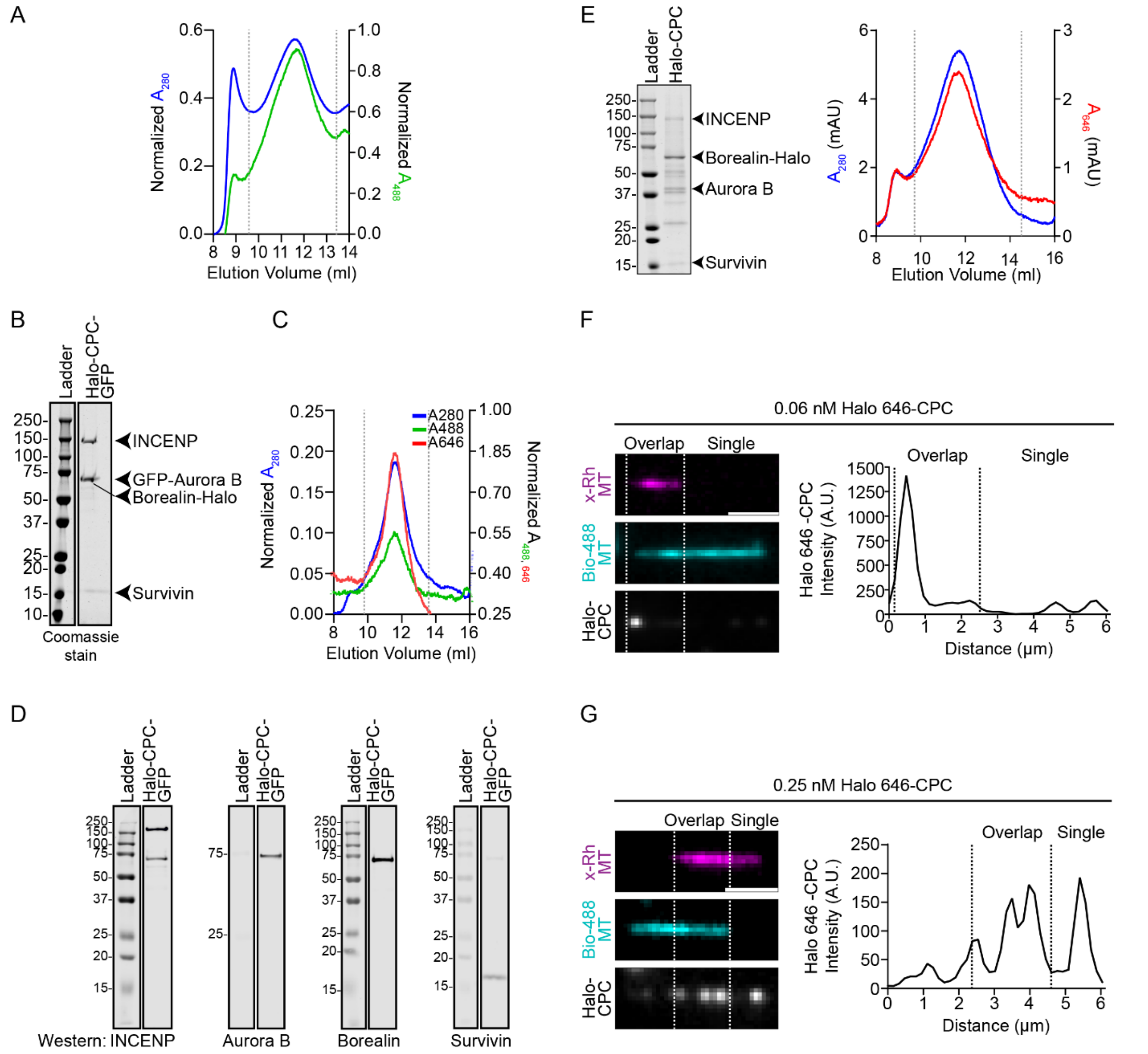

**Figure S1-Related to Figure 1: Purification, characterization, and microtubule-CPC interactions.**

(A) Size exclusion chromatography profile of the CPC-GFP preparation shown in Figure 1B. The graph shows the 280 nm (blue) and 488 nm (green) absorbance traces. Dotted lines show the fractions that were pooled and concentrated.

(B) Coomassie-stained SDS-PAGE 4-20% of purified human Halo-CPC-GFP. Arrowheads show the components of CPC: INCENP, GFP-Aurora B, Borealin-Halo, and Survivin.

(C) Size exclusion chromatography profile of Halo-CPC-GFP. The graph shows the 280 nm (blue), 488 nm (green), and 646 nm (red) absorbance traces. Dotted lines show the fractions that were pooled and concentrated.

(D) Western blots of individual subunits of Halo-CPC-GFP using antibodies specific for INCENP, Aurora B, Borealin, and Survivin.

(E) Coomassie-stained SDS-PAGE 4-20% of Halo 646-CPC. Arrowheads show the components of CPC: INCENP, Aurora B, Borealin-Halo, and Survivin. The graph shows the 280 nm (blue) and 646 nm (red) absorbance traces from size exclusion chromatography of Halo 646-CPC. Dotted lines show the fractions that were pooled and concentrated.

(F-G) The representative images show examples from Figure 1J where, at lower CPC concentrations (0.06 nM and 0.25 nM), high $I_{overlap}/I_{single}$ ratios are observed in some bundles. The three panels show the two microtubules (magenta and cyan) and Halo 646-CPC (grey). The overlap region is indicated by dotted lines. Note: The fluorescence intensity of Halo 646-CPC has been scaled differently in F and G for visibility. The scale bars are 2 µm. The graphs show the intensity profile of Halo 646-CPC along the corresponding microtubule bundles.

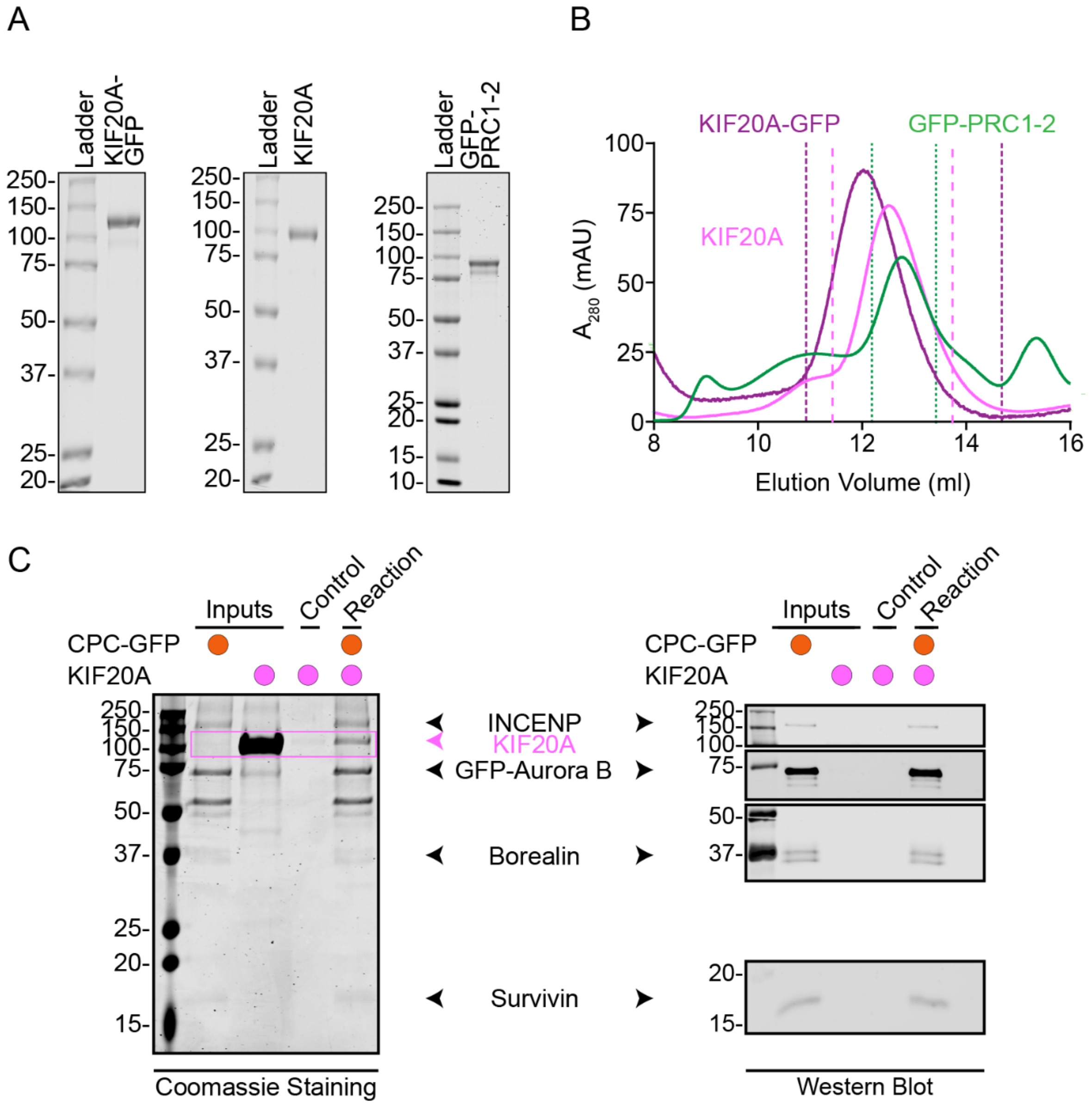


**Figure S2-Related to Figure 2: Purification of recombinant KIF20A and PRC1 isoform 2 and KIF20A-CPC interaction.**

(A) Coomassie-stained SDS-PAGE of purified KIF20A-GFP, KIF20A, and GFP-PRC1 isoform 2.

(B) The graph shows the 280 nm absorbance traces from size exclusion chromatography of KIF20A-GFP (dark magenta), KIF20A (magenta), and GFP-PRC1 isoform 2 (green). Colored dotted lines show the fractions that were pooled and concentrated for each protein.

(C) Coomassie-stained SDS-PAGE (left side panel) from a pull-down assay between CPC-GFP and KIF20A. Arrowheads show INCENP-Flag, GFP-Aurora B, Borealin-Strep, Survivin, and KIF20A. The magenta box indicates the presence of KIF20A. Western blots (right side panel) of individual subunits of CPC-GFP using antibodies specific for Flag, GFP (for Aurora B), Strep (for Borealin), and Survivin.

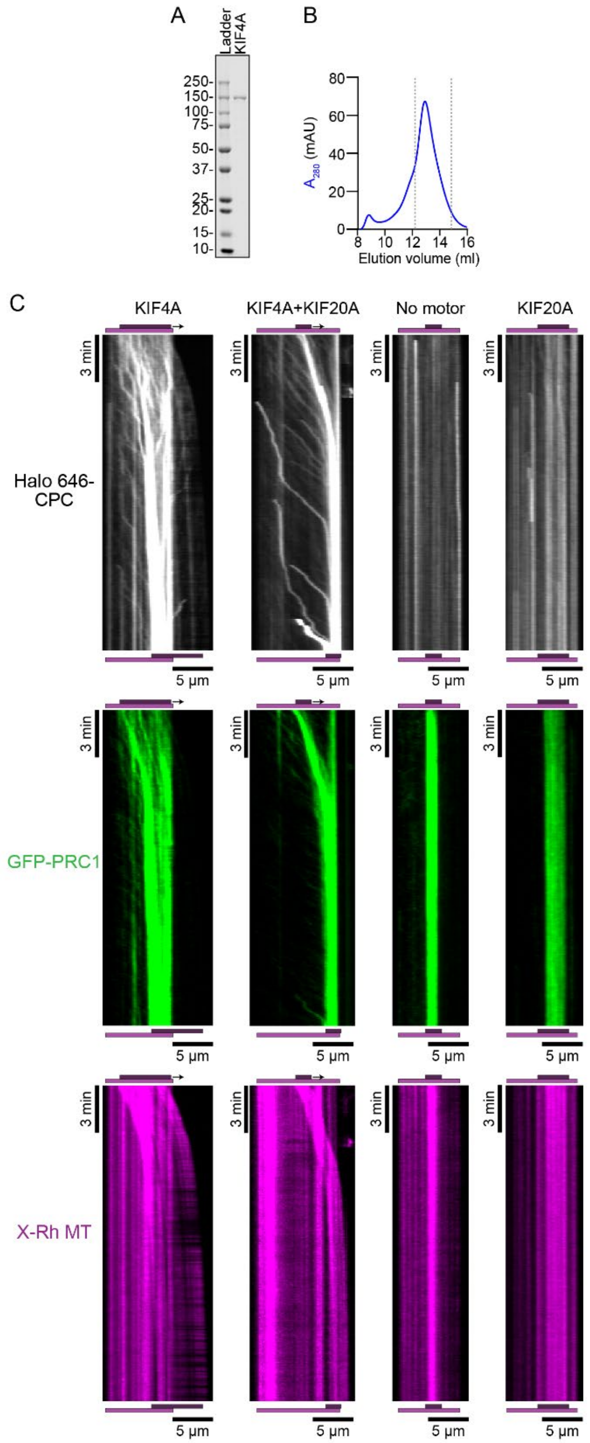


**Figure S3-Related to Figure 3: KIF4A purification and real-time imaging of CPC localization at PRC1-crosslinked microtubule overlaps in the presence of KIF20A and/or KIF4A.**

(A) Coomassie-stained SDS-PAGE 4-20% of purified KIF4A.

(B) The graph shows the 280 nm (blue) absorbance trace from the size exclusion chromatography of KIF4A. Dotted lines show the fractions that were pooled and concentrated.

(C) Representative kymographs from real-time imaging of Halo 646-CPC (2.5 nM, grey) localization on crosslinked microtubules (magenta, lower panel) formed by GFP-PRC1 (green, middle panel) under different conditions: KIF4A (15 nM), KIF4A and KIF20A (15 nM each motor), no motor, and KIF20A (15 nM). The schematics above and below the kymographs indicate the starting and ending positions of the two microtubules. The arrow indicates the direction of movement of the non-biotinylated microtubule relative to the immobilized one (magenta). The scale bars are 3 min and 5 µm.

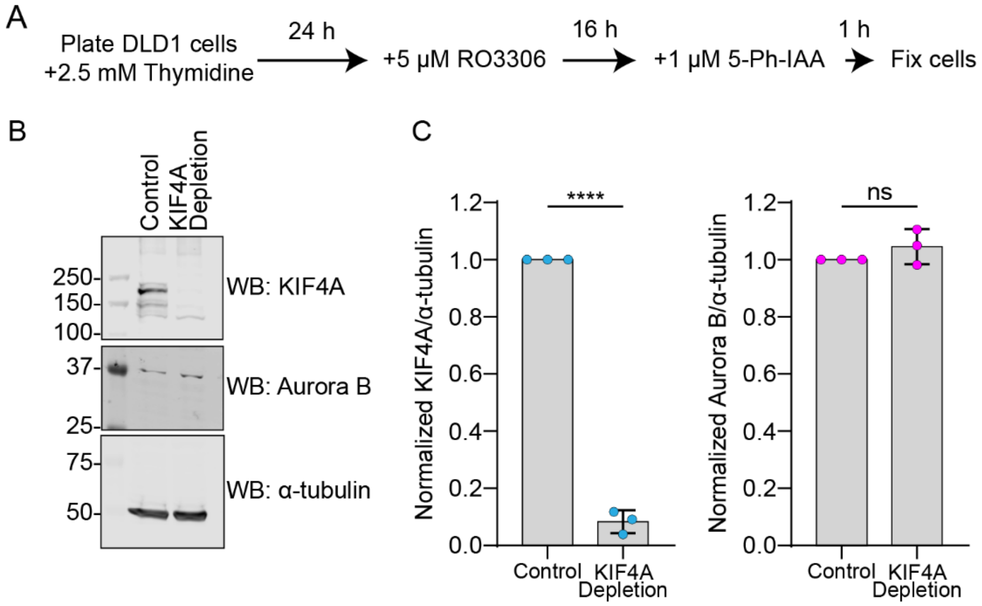


**Figure S4-Related to Figure 4: Effects of KIF4A depletion on Aurora B levels.**

(A) The schematic of the protocol for KIF4A depletion in mitotic cells in DLD1 cells.

(B) Western blot of endogenous KIF4A-SNAP and Aurora B in control and KIF4A-depleted cells. α-tubulin was used as a loading control.

(C) The bar graphs show the normalized KIF4A-to-α-tubulin and Aurora B-to-α-tubulin intensities in control and KIF4A-depleted cells from the western blot experiments in B (N=3).

**Movie captions**

Movie 1: Example TIRFM videos of Halo 646-CPC (2.5 nM) in the presence of (A) KIF4A (15 nM), (B) KIF4A and KIF20A (15 nM each motor), (C) no motor, and (D) KIF20A (15 nM)-related to Figure 3 and S3. Time is shown as minutes:seconds. The scale bars are 5 μm.
